# spinDrop: a droplet microfluidic platform to maximise single-cell sequencing information content

**DOI:** 10.1101/2023.01.12.523500

**Authors:** Joachim De Jonghe, Tomasz S. Kaminski, David B. Morse, Marcin Tabaka, Anna L. Ellermann, Timo N. Kohler, Gianluca Amadei, Charlotte Handford, Gregory M. Findlay, Magdalena Zernicka-Goetz, Sarah A. Teichmann, Florian Hollfelder

## Abstract

Droplet microfluidic methods have massively increased the throughput of single-cell sequencing campaigns. The benefit of scale-up is, however, accompanied by increased background noise when processing challenging samples and the overall RNA capture efficiency is lower. These drawbacks stem from the lack of strategies to enrich for high-quality material or specific cell types at the moment of cell encapsulation and the absence of implementable multi-step enzymatic processes that increase capture. Here we alleviate both bottlenecks using fluorescence-activated droplet sorting to enrich for droplets that contain single viable cells, intact nuclei, fixed cells or target cell types and use reagent addition to droplets by picoinjection to perform multi-step lysis and reverse transcription. Our methodology increases gene detection rates fivefold, while reducing background noise by up to half. We harness these unique properties to deliver a high-quality molecular atlas of mouse brain development, despite starting with highly damaged input material, and provide an atlas of nascent RNA transcription during mouse organogenesis. Our method is broadly applicable to other droplet-based workflows to deliver sensitive and accurate single-cell profiling at a reduced cost.

## Introduction

Droplet microfluidic methods have fundamentally transformed the field of single-cell RNA-sequencing (scRNA-seq) by increasing the number of cells that can be profiled in a single experiment by more than an order of magnitude compared to plate-based assays^1^. These technological advances have propelled the field into the age of molecular atlases that aim to resolve the full spectrum of cellular heterogeneity across entire organs^2–8^ or organisms^9,10^. Although combinatorial indexing methods, such as sci-RNA-seq3^11^ surpass droplet microfluidic methods in terms of throughput^1^, most atlases to date have been generated using the commercial 10x Chromium platform^3^, which can be explained by earlier adoption and extensive standardisation of commercial kits and analysis software^12^. Although the popularity of droplet-based approaches for single-cell profiling is evident, some fundamental challenges associated with the methodology remain to be resolved.

The vast majority of single-cell RNA sequencing campaigns will aim to maximise the number of cells profiled and the number of unique cDNA molecules that can be confidently detected per cell, sometimes referred to as sensitivity^13^, to yield statistical power to downstream analyses such as differential gene expression analysis. However, there is currently a trade-off between cost of library preparation and gene detection rates per cell. Although the commercial 10x Chromium^12^ outclasses open-source protocols such as inDrop^14^ and Drop-seq^15^ in terms of sensitivity (2.5- and 1.2-fold higher gene detection rates, respectively)^16^, the associated library preparation cost per cell remains prohibitive for large-scale profiling (twofold higher than respective open-source methods). Therefore, molecular atlasing experiments would hugely benefit from a method with reduced cost per cell while maintaining high sensitivity to derive meaningful biological conclusions.

Furthermore, the quality of the data derived from droplet-based methodologies can suffer from artefacts that may confound data analysis: RNA released from lysed cells indiscriminately enters into droplet compartments and generates a contaminating readout that is not cell-specific anymore, contributing both to cost and compromising data interpretation^17–19^. Cell debris or damaged cells can also be captured in those experiments, further complicating the identification of live cells in a dataset. Although experimental safeguards can be implemented to mitigate these effects, such as pre-sorting of live cells using fluorescence or magnetic-associated cell sorting, they do not guarantee viability at the moment of encapsulation. On the contrary, shear stress during flow cytometry or long processing times may lead to altered transcriptomes^20^ or cell death^21^. Live cell enrichment procedures typically require large amounts of input material^22^ and do not remove contaminating sequences, such as primer dimers and concatemers, generated from empty droplets. Similarly, bioinformatic tools (EmptyDrops^18^, SoupX^17^, emptyNN^23^ and DropletQC^19^) have been employed to remove confounding effects from empty droplets and/or low-quality material. However, the performance of these approaches depends on sample quality and the cell types profiled. Finally, cell multiplets resulting from cell co-encapsulation or aggregation may further compromise data analysis by generating artificial cell populations. Although tools have been developed for their identification during data processing^24,25^, they mostly resolve heterotypic multiplets (i.e. multiplets containing cells from different cell types), and do not remove the burden of sequencing costs associated with these artefacts. To this end, a generalizable methodology to extract droplets containing single viable cells, intact nuclei or specific cell types from the pool of empty droplets, droplets containing cell debris, multiplets or unwanted cell types would reduce cost and remove confounding artefacts found in droplet microfluidic datasets.

To alleviate the aforementioned bottlenecks, we have developed **s**orting **p**icoinjection **inDrop** (spinDrop), a scalable droplet microfluidic method that delivers highly-sensitive 3’ mRNA sequencing of single viable cells, intact nuclei, paraformaldehyde-fixed samples or specific cell types at a reduced cost. The protocol first employs fluorescence-activated droplet sorting (FADS)^26^ to exclusively extract target material followed by a picoinjection step^27^ of an improved reverse transcription formulation. We demonstrate five-fold higher gene detection rates compared to inDrop^14^ to match the resolution obtained with the 10x Chromium^12^, while significantly reducing the noise linked to empty droplets and poor quality cells. We demonstrate the utility of our workflow by profiling mouse brain development using a damaged sample as input while maintaining high data output quality. The multi-step capabilities of our workflow to power new high-throughput ‘-omics’ applications were demonstrated by profiling nascent RNA transcription during mouse organogenesis at E8.5 using 5EU-seq implemented in droplets.

## Results

### Overview of the spinDrop workflow

The spinDrop microfluidic workflow aims to generate highly-sensitive single-cell RNA-seq libraries from small quantities of input material with minimal contamination from damaged cells and empty droplets. To achieve this, a multi-step droplet microfluidic workflow was established (Figure 1A). First, a strategy was devised to alleviate the current bottlenecks of pre-sorting for viability using FACS or MACS, which necessitate long processing times, large input materials and may damage the input material further due to mechanical shearing. To this end, we implemented in-line FADS^26^ to enrich for droplets that match a sorting *criterion* (e.g. reporting on cell viability) after cell co-encapsulation with barcoded microgels in water-in-oil emulsions. Cells are first stained using a viability dye (Calcein-acetoxymethyl (AM)). For sorting of fixed cells or intact nuclei, a DNA stain can be employed (Vybrant Green) instead. For cell-type specific enrichment, fluorescently labelled antibodies targeting cell surface markers are used. The cells are then channelled in a flow-focusing device in conjunction with barcoded polyacrylamide microgels (with an inDrop v3 barcoding scheme^28,29^) and a lysis mixture (Figure 1B). Single-cells are hereby co-encapsulated with the microgels in water-in-oil emulsions and can be sorted further down in the microfluidic device. Cells trapped in a droplet are excited by an adjacent blue-laser optical emission fibre. If cells are viable at the moment of encapsulation, the acetoxymethyl group linked to the Calcein fluorophore (in the case of live cell sorting) is released by intracellular esterases^30^, enabling green fluorescence emission via blue-light excitation. Fluorescence is then collected in a second optical fibre, which relays the information to a field-programmable gate array which controls an amplifier and square signal AC generator (Supplementary Figure 1). If the detected levels of fluorescence exceed the negative background signal generated from empty droplets or droplets containing damaged cells, an electrical pulse is triggered via the activation of electrodes filled with a 5 M NaCl solution^31^. The activation of electrodes charges the droplets via dielectrophoresis and diverts them from the lower hydrodynamic resistance channel (negative channel) to pull them towards the positive channel (Figure 1C, Extended Data Figure 1A, 2A-B, Supplementary video 1) for downstream processing.

**Figure 1.**
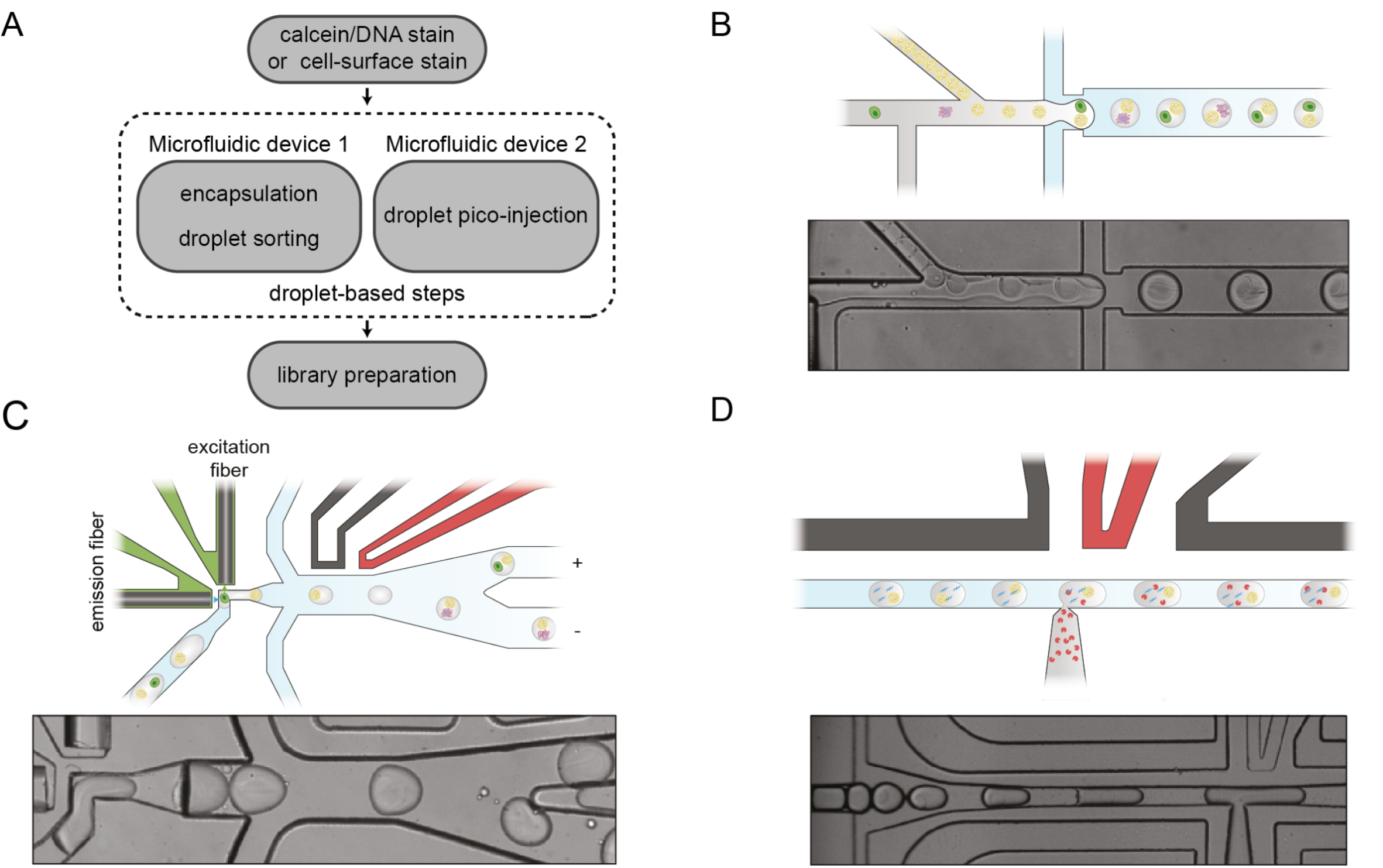
Overview of the modular droplet microfluidic workflow for spinDrop (‘**s**orting and **p**icoinjection **inDrop**’). A) Schematic of the different steps to generate sequencing-ready libraries. First intact whole cells are made detectable with a Calcein-AM viability stain, cell-surface marker stain, and intact nuclei using a DNA staining dye (such as the Vybrant DNA stain). Then the cells are co-encapsulated with barcoded polyacrylamide beads, and viable cells, intact nuclei or specific cell types are enriched after encapsulation using in-line fluorescence-activated droplet sorting (FADS), hereby discarding empty droplets or droplets containing damaged material. The droplets containing the viable material are then lysed and heat-treated, and further re-injected in a second microfluidic device, which will inject the reverse transcriptase mix using coalescence-activated picoinjection. B) Schematic of the cell/nuclei and polyacrylamide bead droplet co-encapsulation process in the first microfluidic device (*top*). Brightfield image of the co-encapsulation process (*bottom*). C) Schematic of FADS-sorting of droplets containing viable cells or intact nuclei from the pool of empty droplets containing damaged or unwanted material (*top*). Brightfield image of the in-line sorting process (*bottom*). D) Schematic of the coalescence-activated droplet picoinjection of reverse transcriptase mix (*top*). Brightfield image of the picoinjection process (*bottom*).

Second, the sensitivity bottleneck of open-source platforms, in particular inDrop^14^ upon which this workflow is based on, was addressed using multi-step enzymatic and incubation processing. Currently, droplet microfluidic single-cell protocols rely on a single enzymatic treatment at fixed temperatures to perform reverse transcription^12,14,15,32^due to incompatibility of lysis via proteolysis and reverse transcription enzymatic reaction. This limitation prevents enhanced RNA capture rates observed in plate-based assays, which, for example, use proteinase K and heat denaturation during the lysis step to increase denaturation of the nuclear envelope, release nucleic acids from protein complexes and unfold secondary structures to boost RNA capture^33–37^. We previously described a droplet microfluidic methodology that enables multi-step reactions for single-cell sequencing to be carried out in droplets^37^, employing a previously described microfluidic design termed ‘picoinjector’^27^(Extended Data Figure 1B-C). We therefore sought to use this method to decouple cell lysis from reverse transcription (RT) by injecting a RT reaction mixture optimised for 3’ mRNA capture consecutively to cell lysis, to match workflows from the more sensitive plate-based assays. To achieve this, water-in-oil emulsions containing the sorted single-cell lysate are loaded into the picoinjector and spaced using fluorosurfactant oil. When approaching a junction with an electrode, the droplet interface is dielectrophoretically disrupted and the pressurised flow of mixture is injected into the incoming droplet (Figure 1D, Supplementary video 2). The droplets can then be further collected and incubated for reverse transcription before de-emulsification and downstream library preparation, following the inDrop protocol^14^.

### FADS reduces background noise and increases cell loading in droplets beyond Poisson distribution

To characterise the capabilities of FADS to enrich for encapsulated single cells, HEK293T cells were stained with Calcein-AM and sorted using a threshold that separated the background signal from the cellular fluorescence signal (Figure 2A, Extended Data Figure 2C) without the addition of a cell lysis mix in the final emulsion. The resulting sorted droplets were analysed on a fluorescence microscope, showing a stark enrichment (96.1%, n= 51) for droplets containing single viable cells (Figure 2B). To further quantify the potential of our system to extract single viable cells from a challenging sample containing a large proportion of damaged cells, the input population was modified to incorporate a 1:1 ratio of dead and alive HEK293T cells treated with a dual live/dead stain (Calcein-AM and ethidium homodimer-1, respectively). To induce cell death, a concentration of 0.25% (w/v) IGEPAL CA-630 was added to half of the HEK293T and incubated on ice for 15 minutes. Sorting of a 1:1 mixed population of dead and living cells showed a marked 19-fold enrichment for viable cells from the pool of droplets, with 84.8% of the droplets containing a single viable cell, which surpasses the predicted value of 4.52% without sorting (Figure 2C). On the other hand, the remainder of the droplets collected in the negative channel mostly contained 93% of empty droplets, 3.5% dead cells and 3.1% living cells (3.1%) (Figure 2C). To assess whether discarding cellular multiplets as a result of stochastic co-encapsulation is possible, an upper threshold on the fluorescence signal (3V) was applied to a sample with a fivefold higher Poisson loading (defined by λ value which is the mean number of cells per droplet) of 1:1 mixed dead and living cells, to exclude brighter droplets that may contain multiple cells. Under these conditions, 30.3% of sampled droplets are predicted to contain a single cell (compared to 9.0% at λ=0.1), which represents a threefold increase in processing throughput and may assist large-scale sampling endeavours, but comes at the cost of higher multiplet rates. At λ=0.5, 77.2% of the sorted droplets contained a single viable cell after sorting, which largely outclasses the predicted values (Figure 2C). While viable cells can be detected, residual dead cells that do not emit fluorescence can be co-encapsulated with viable cells which cannot discriminated against using FADS. The proportion of droplets containing one dead and one living cell in the sorted population was 5.6%, slightly higher than the predicted value of 3.8%. This limit is due to stochastic co-encapsulation of invisible cells and remains when isolating single viable cells at higher loading concentrations for input samples with low variability.

**Figure 2.**
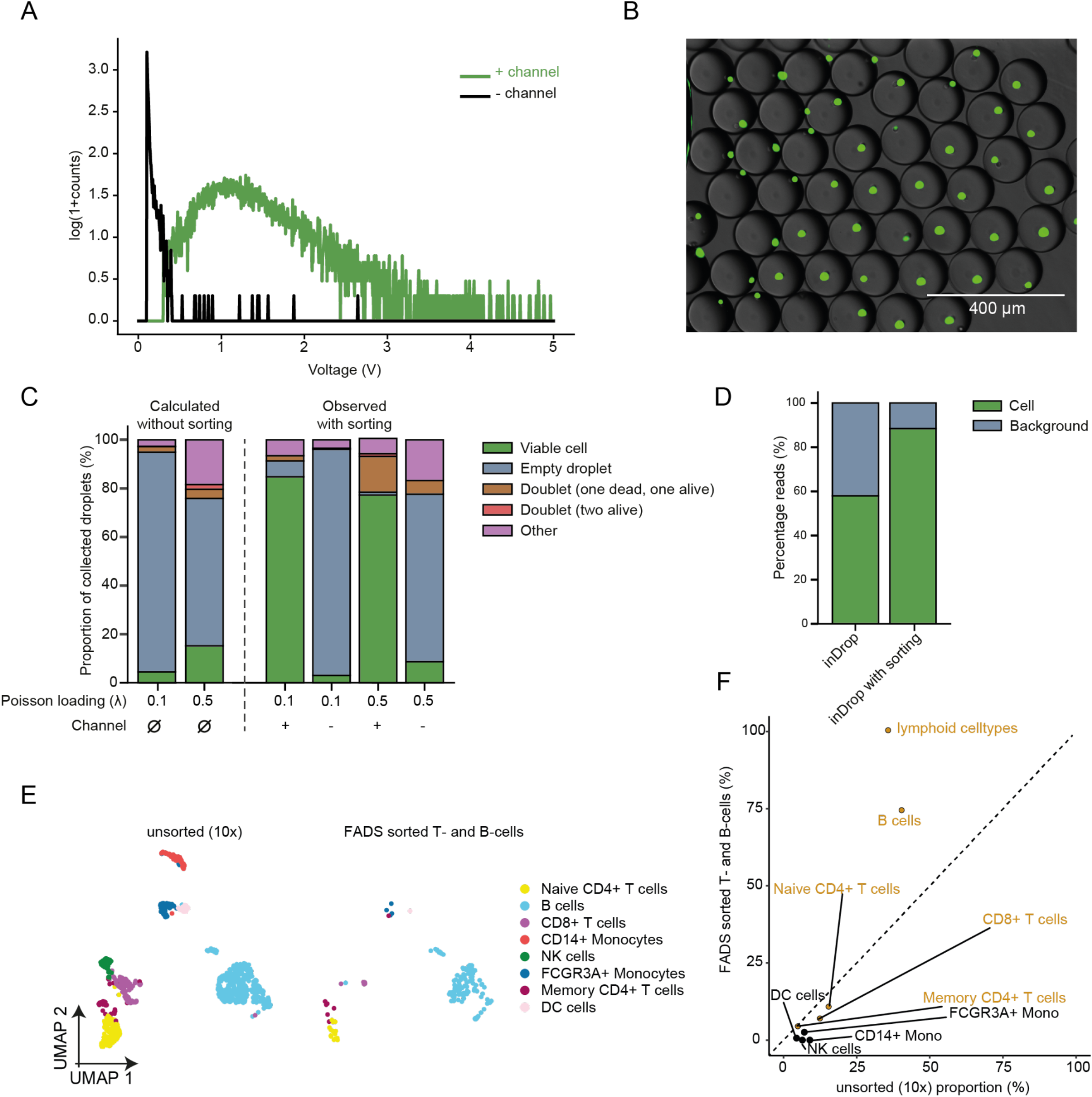
Sorting of viable cells using FADS decreases background noise from cell-free RNA and dead cells. A) Measured signal of HEK293T cells stained with Calcein-AM using a detection fibre and photomultiplier-tube, with thresholding on the background population (black) and the droplets with an encapsulated viable cell (green). B) Overlay of fluorescence and brightfield images of the positive sorted population of droplets, showing the accurate single compartmentalisation of viable cells. Scale bar: 400 μm. C) Sorting statistics using a dual live/dead stain on a 1:1 dead/alive population of HEK293T cells. Poissonian loading stands for a cell loading of λ = 0.1, super-loading stands for a cell loading of λ = 0.5. (+) indicates counting on the sorted droplet, (-) indicates counting on the unsorted droplets. The predicted values without sorting are on the left side of the dashed line, the observed values are on the right side of the dashed line. D) Percentage of reads that are mapped to background and cell barcodes for inDrop and inDrop with sorting, determined using the filtering statistics from the zUMIs pipeline^43^. E) UMAP dimensional reduction plot showing the shared embedding for mouse PBMCs sequenced using the 10x v2 protocol and sequenced after on-chip FADS sorting using CD45R, CD19 and IgM as cell-surface specific fluorescent antibodies. F) Proportional distribution of each cell type in the dataset in E), illustrating an enrichment for lymphoid B- and T-cells in the FADS sorted dataset. The dotted line represents equal proportions across both datasets.

To further determine if discarding empty droplets from the analysis would lead to a lower fraction of background reads generated by empty droplets, a species-mixing experiment with human HEK293T and mouse embryonic stem cells (mESCs) was performed (under inDrop reaction conditions), with and without sorting. The proportion of reads matching to a cell barcode through the computation of a coverage inflection point was 58.0% for inDrop without sorting, similarly to previously reported values^14^ and 88.4% for inDrop with sorting (Figure 2D), documenting a clear gain in cellular read coverage for cultured cells that translates into lower sequencing costs and higher accuracy for data interpretation.

To further test the ability of our system to enrich for specific cell types using FADS, mouse peripheral blood mononuclear cells (PBMCs) were stained using phycoerythrin (PE) labelled antibodies specific to CD19, CD45R and IgM and processed for sequencing. Projecting the dataset on a UMAP embedding containing an unsorted reference dataset generated using the 10x Chromium revealed a marked depletion of myeloid cell types (Figure 2E, Extended Data Figure 2E), and an overall enrichment of 1.8-fold in B-cell content. Monocytes, such as CD14+ monocytes were entirely depleted, whereas they accounted for 9% of the population without enrichment (Figure 2F).

These findings illustrate the benefits associated with in-line droplet sorting using FADS: to reduce cost and decrease potential confounding effects from empty droplets and degraded input material. Additionally, cell-type specific enrichment may further decrease cost and enable higher coverage of rare cell populations, in applications where FACS is incompatible, because of low cell numbers or viability.

### Improved lysis and reverse transcription reaction conditions increase RNA capture

We next sought to improve RNA capture efficiency in our assay, which is a current bottleneck in open-source workflows such as inDrop^14^. In existing droplet-based single-cell transcriptomics approaches, including the 10x Chromium system, the content of the droplets cannot be modulated after encapsulation. This precludes implementation of efficient cell lysis before reverse transcription, as best-in-class plate-based assays use heat denaturation and proteinase K treatments to increase RNA yields^33,34^, which are fundamentally incompatible with reverse transcription. We therefore sought to decouple both steps by performing lysis at the encapsulation stage followed by reverse transcription after picoinjection of the droplets containing the cell lysate in a second instance. We tested several reaction conditions using HEK293T cells: 1) the addition of 7.5% (w/v) PEG 8000 as a crowding agent to increase sensitivity (as in the plate-based mcSCRB-seq protocol^34^); 2), the use of proteinase K and heat denaturation in addition to the non-ionic detergent IGEPAL CA-630 used in the inDrop protocol; 3) different reverse transcriptases, mainly Maxima H- and Superscript III RTs used in mcSCRB-seq^34^ and Smart-seq3^33^, respectively. Downsampling the dataset to 20,000 reads per cell across conditions revealed a gene detection rate of 1,016 genes for the original inDrop reaction conditions, in line with previous findings^14,16^. Adding 7.5% (w/v) PEG8000 and decoupling lysis from RT resulted in a median of 3,384 (with PEG) and 3,403 (without PEG) detected genes per cell, demonstrating that molecular crowding does not yield higher sensitivity in droplet formats. However, performing lysis with a proteinase K and heat denaturation treatment increased gene detection rates more than threefold compared to inDrop, leading to an overall detection of 4,005 genes using the Maxima reverse transcriptase and 4,926 using Superscript III (Figure 3A). Because the last conditions demonstrated superior performance, the remainder of the manuscript will utilise this set of reaction conditions (dubbed ‘spinDrop’). The gain in the number of genes detected per cell is accompanied by higher detection-levels of intronic reads, which amount to 36.8% for spinDrop (Extended Data Figure 3A) compared to 31.8% for inDrop, which may benefit RNA velocity analyses^38^.

**Figure 3.**
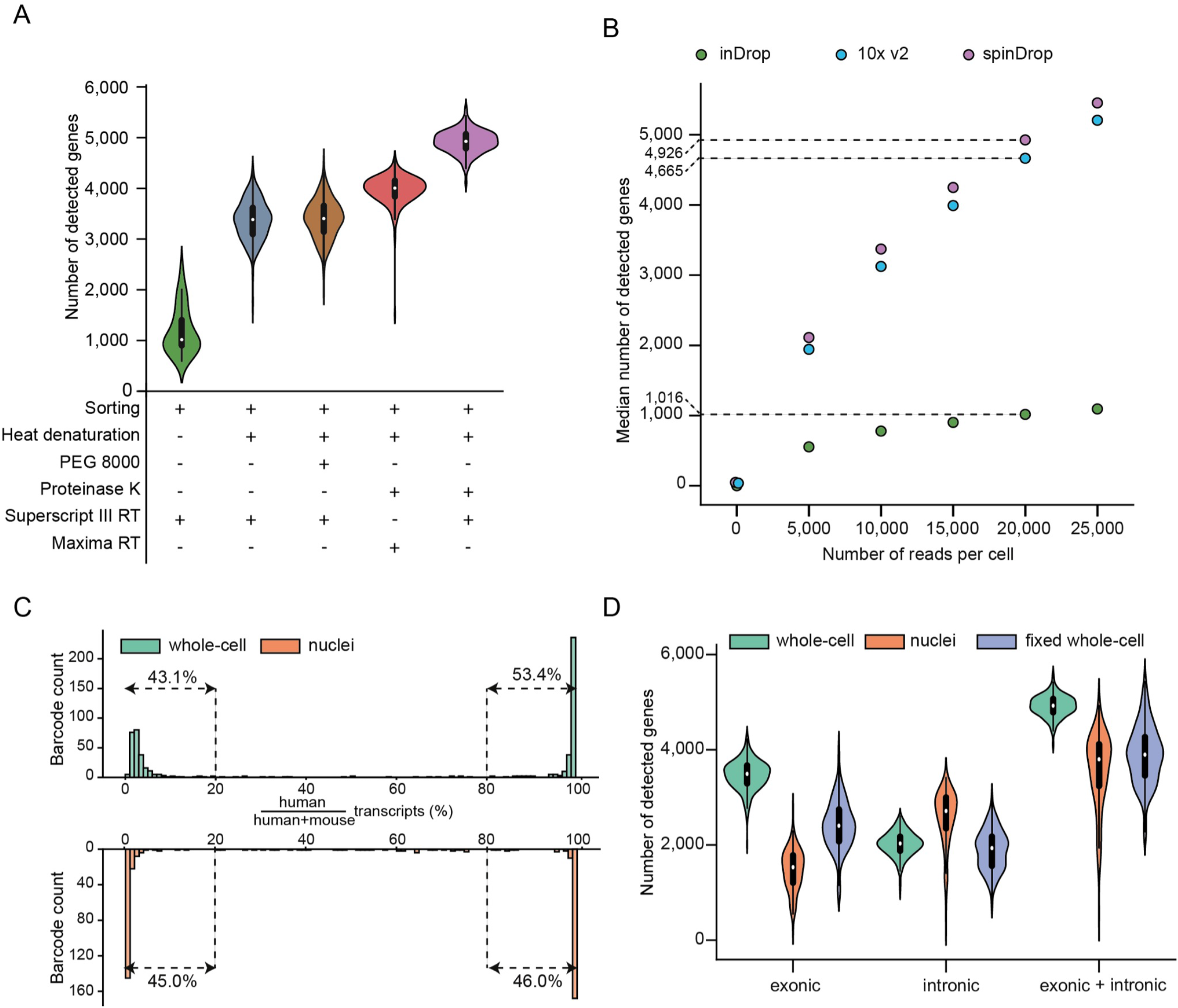
Improved lysis procedures and picoinjection of reverse transcriptase mixture increase RNA capture efficiency. A) Gene detection rates for different reaction conditions using Calcein-AM stained HEK293T cells. Each dataset was downsampled to 20,000 reads per cell to allow for direct comparison between reaction conditions. The purple colour delineates the conditions for sorting picoinjection inDrop (spinDrop) used in the remainder of the manuscript. B) Gene detection downsampling curves comparing the median number of genes detected for HEK293T cells with the 10x Chromium v2, inDrop and spinDrop reaction conditions. The biological read length for each sample was cut to 61 bp to allow for a direct comparison between all datasets. The dashed line indicates median gene detection rates at 20,000 reads per cell. C) Species mixing barcode proportional counts using mouse embryonic stem cells and HEK293T cells (green) or nuclei (orange) as an input for the spinDrop protocol. D) Intronic and exonic UMI counts for HEK293T cells either prepared as whole cells (green), nuclei (orange) or fixed whole cells (purple), illustrating high capture across different input formats.

These improvements were then measured against the most sensitive droplet microfluidic platform^12^, the 10x Chromium, by downsampling read cellular coverage from 5,000 to 25,000 reads per cell to evaluate gene detection saturation. The downsampling curves demonstrated slightly lower sensitivity for the 10x Chromium platform compared to spinDrop, with a median of 4,665 genes detected using 10x Chromium at 20,000 reads per cell (Figure 3B). A Wilcoxon rank sum test further established the core differences in the genes detected between the 10x Chromium and the spinDrop methodology. The analysis revealed a total of 690 genes were significantly and robustly detected throughout the dataset (absolute values for log2 fold change > 1 and Bonferroni adjusted p-values <10^−5^). Further annotation of the genes by biotypes showed an enrichment for non-coding RNAs and pseudogenes for spinDrop and some protein-coding genes in the 10x Chromium dataset (Extended Data Figure 3C, Supplementary table 1). From the list of significantly differentially expressed genes, spinDrop showed an enrichment of 291 genes which, classified proportionally per biotype, were: 1) 2.1% lncRNAs, 2) 26.4% processed pseudogenes,

63.9% protein-coding, 4) 2.7% transcribed processed pseudogenes and 5) 2.7% unprocessed pseudogenes. The 10x dataset, on the other hand, had 399 genes that were upregulated which, classified proportionally per biotype, were: 1) 99% protein coding genes and 2) 1% of lncRNAs. This illustrates some of the core differences between both methods, likely owing to differences in library preparation methods, such as the cDNA amplification process and lysis method. For example, spinDrop uses *in vitro* transcription for cDNA amplification and 10x Chromium uses template-switching PCR-based for cDNA amplification, which has been reported to yield different gene capture^39^. Another fundamental difference relies on the release of nuclei-localised RNA molecules such as lncRNAs, which could be found at higher levels in the spinDrop methodology, likely owing to more efficient lysis of the nuclear envelope due to the proteinase K and heat denaturation steps. Technology-specific preferential enrichment of gene classes may further inform mechanisms. For example, some of the enriched protein coding genes for the spinDrop method relate to Gene Ontology terms GO:0048024 (regulation of mRNA splicing, via spliceosome, FDR 2.0 e^-8^) and GO:0043484 (regulation of RNA splicing, FDR 7.6 e^-11^), hinting towards better definition of the splicing machinery in this methodology (Extended Data Figure 3D). In addition, the spinDrop method enables capture of molecular spike-ins which is prohibitively expensive in traditional droplet set-ups due to the overwhelming majority of droplets being empty. For example ERCC^40^ molecules, which can be utilised to assist data normalisation, were readily captured using spinDrop without the sequencing cost burden associated with empty droplet capture (Extended Data Figure 3E).

To verify that the spinDrop method reliably compartmentalised single-cells and preserved the compartment throughout the picoinjection step, a species mixing assay comprising mESCs and human HEK293T cells was performed in both cellular and nuclei formats (Figure 3C). The analysis revealed a doublet rate of 2.9% for the cellular sample (Extended Data Figure 4A) and 6.1% for the nuclei sample (Extended Data Figure 4B), illustrating a low proportion of doublets and droplet merging throughout the microfluidic process. Finally, we used spinDrop to process paraformaldehyde-fixed HEK293T cells. Cell fixation is the method of choice for archiving clinical samples^41^ and may position our methodology as uniquely suited to interrogate previously inaccessible sample types from repositories. Contrasting all three formulations showed that nuclei had higher numbers of reads mapping to intronic regions (median of 2,714 genes) compared to exonic regions (median of 1,533 genes). This distribution was reversed for the whole cell samples, which detected a median of 2,030 genes with reads mapping to introns and 3,493 genes with reads mapping to exons. The sample with fixed cells displayed slightly lower gene detection rates, with 1,934 genes with reads mapping to introns and 2,404 genes with reads mapping to exons (Figure 3D).

Therefore, adding a picoinjection module to the workflow enabled a fivefold gain in gene detection rates compared to inDrop (4,926 vs 1,016, Figure 3A), with increased RNA biotype diversity and intron detection. The sensitivity is conserved throughout different input formats, which makes spinDrop a versatile and sensitive platform for scRNA-seq.

### Profiling of the embryonic mouse brain using a highly damaged sample input

We further investigated whether the sorting feature of the spinDrop platform could be leveraged to improve the quality of biological samples with low viability. Such samples contain a larger amount of degraded or disintegrated cells, or cells in stressed states that might not reflect a physiologically relevant transcriptional programme^21,42^. To this end, we dissected and dissociated the brain of mouse embryos at developmental stage E10.5 and left it at room temperature for 3 hours to decrease viability (Figure 4A), which amounted to 36.6% after Calcein-AM staining and counting on a haemocytometer. We then processed the cells with and without sorting to demonstrate the utility of our workflow to tackle complex samples. Ranked read coverages per barcode (knee plots) were generated for both samples using zUMIs^43^, which underlined a clear inflexion point for the sample where pre-sorting was used, which contrasted strongly with the unsorted sample which had a linear trend (Figure 4B). This rebalancing towards *bona fide* cells with increased coverage suggests an enrichment for high-quality material using FADS sorting.

**Figure 4.**
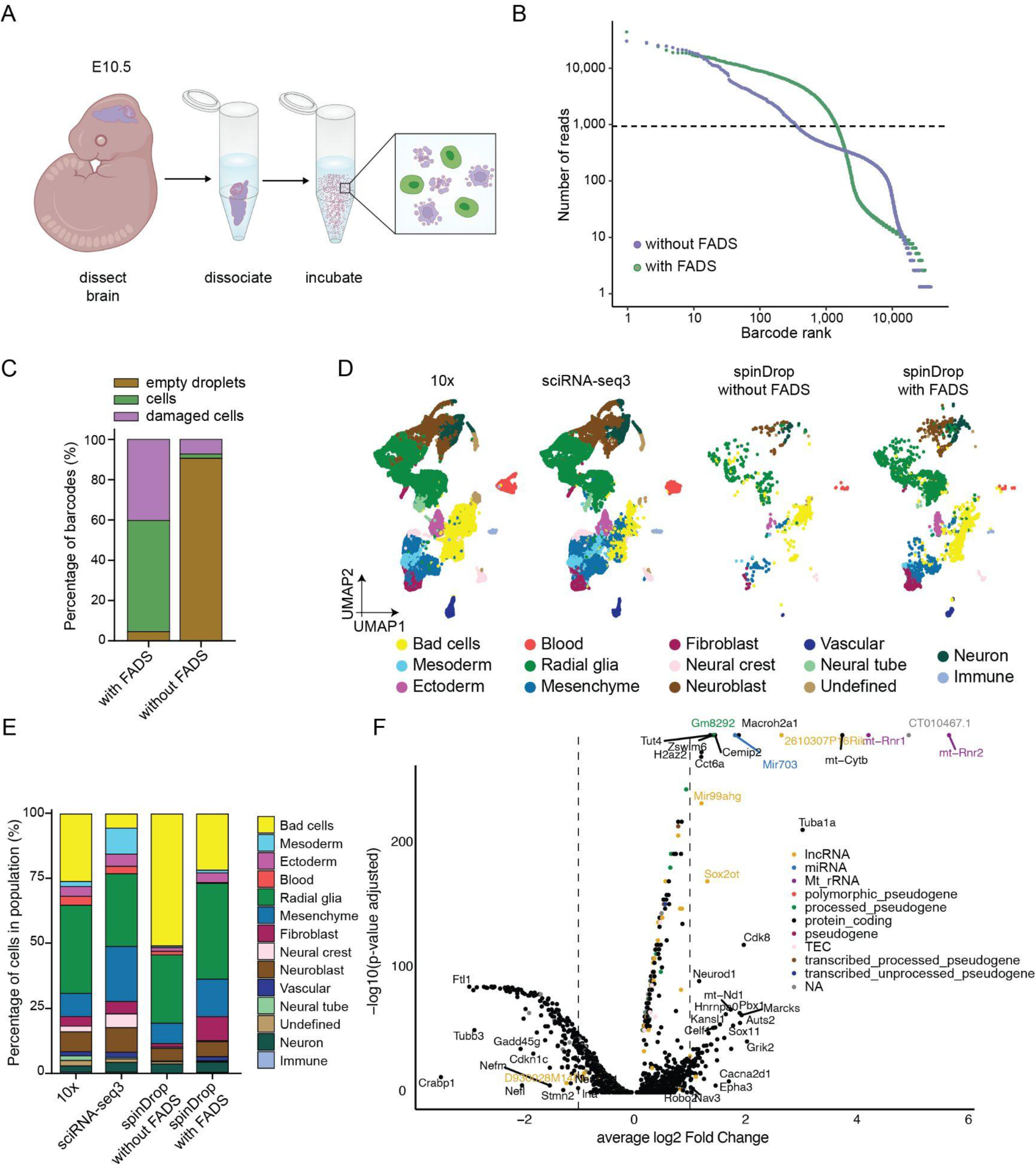
Transcriptional atlas of embryonic mouse brains at E10.5 generated using the spinDrop methodology A) Schematic of the cell recovery process and staining. The cells are stained using Calcein-AM for separating live cells (green) from dead cells (purple). The cells were left in PBS at room temperature for three hours after dissociation to increase cell death rates. B) Barcode rank knee plot for the mouse brain dataset using in-line sorting to extract live cells (green, with FADS), or without sorting to denote results using standard droplet microfluidic scRNA-seq (purple, without FADS). The sorted population shows a barcode inflection point at ∼1,000 barcodes, contrary to the sorted population, illustrating lower background from dead cells and cell-free RNA. C) Results from barcode quality control inspection using the DropletQC tool for both the droplets with FADS sorting and without FADS sorting. The sorted population shows a significantly lower proportion of empty droplets and damaged cells compared to the unsorted population. D) UMAP dimensional reduction on an equivalent 10x, sciRNA-seq3 and spinDrop dataset (with and without sorting), cells labelled by major cell type. n= 30,244 cells for the 10x dataset, 20,000 cells for the sciRNA-seq3 dataset, 2,472 cells for spinDrop with FADS sorting and 802 cells for spinDrop without FADS sorting. E) Histogram representing the proportion of cells per major cell-types for each technology. F) Volcano plot representing the differentially expressed genes for the Neuroblast cluster between 10x (negative log2 fold change values) and spinDrop with sorting (positive log2 fold change values).

The datasets were then further investigated using DropletQC^19^ to identify empty droplet barcodes as well as dead cells, computed using nuclear fractions (ratio of unspliced to spliced reads) on both sorted and unsorted samples. The analysis revealed that 90.7% and 1.2% of the barcodes were determined as empty droplets and damaged cells in the unsorted population (Extended Data Figure 4E). By contrast, these proportions were 4.6% and 40.2% respectively with FADS sorting (Extended Data Figure 4F), illustrating efficient elimination of spurious transcripts by discarding empty droplets using FADS (Figure 4C). To verify if the datasets generated would be amenable to transcriptional atlasing of heterogeneous cell types, the spinDrop datasets were integrated with mouse embryo references generated using the 10x v1^44^ (E11) and sci-RNA-seq3^11^ (E10.5, downsampled to 20,000 cells to match other sample sizes) methods as references. To this end, Seurat v3 was utilised to compute integration anchors and generate a shared embedding^45^ between all datasets, and further computed Pearson correlation coefficients between the clusters in the shared embedding and the annotation from La Manno *et al*.^*44*^ were used to transfer labels to all remaining datasets (see Methods) (Figure 4D). The cell type distributions after label transfer delineated a decrease in the proportion of cells in the low complexity cluster (“Bad cells” in yellow in Figure 4D) when applying FADS sorting to the input population, from 50.9% without sorting to 21.7% with sorting and 26.1% with the 10x dataset (Figure 4E). To further investigate if the dataset generated using our methodology could improve marker delineation in the dataset, a Wilcoxon rank sum test was used to compute differential marker analysis between the 10x and spinDrop dataset (with sorting) on the Neuroblast population (Supplementary table 2). The analysis revealed an enrichment for core cortical maturation markers, such as *Sox11*^*46*^, *Sox2ot*^*47*^ and *Neurod1*^*48*^ using the spinDrop methodology.

These findings underline the capabilities of our workflow to capture the spectrum of heterogeneity in a damaged input sample by removing damaged cells and empty droplets from the sampled population.

### High-throughput nascent RNA sequencing using spinDrop

SpinDrop is a multi-step method enabling complex RNA capture protocols that may not be attainable using single-step standard droplet tools because of temperature, enzymatic and buffer incompatibilities^12,14,15^. For example, performing reverse cross-linking in droplets is currently prohibited in droplets as mentioned previously. This bottleneck prevents the scaling-up of methods that use fixation as a means to broaden transcriptome read-outs such as metabolic labelling methods like 5EU-seq^49^. We therefore applied the methodology to uncover nascent RNA transcription during mouse organogenesis at high-throughput. To achieve this, mouse embryos at E8.5 were dissected and cultured *in vitro* for 3 hours with the 5EU (5-Ethynyl Uridine) analog, then dissociated, fixed and subjected to click chemistry to add biotin groups to the analog incorporated in the nascent transcript. The fixed cells were then processed using spinDrop for reverse cross-linking and reverse transcription, without sorting, and the nascent transcripts were pulled-down using streptavidin coated beads before performing library preparation on both nascent and non-nascent transcripts (Figure 5A). Nascent and non-nascent RNA molecules for each cell were then linked bioinformatically. The nascent RNA libraries displayed a higher proportion of UMIs mapping to intronic regions of transcripts than the non-nascent library (43.4% versus 21.4%, ratios of median UMI counts), confirming higher capture rates for nascent transcripts (Figure 5B, Extended Data Figure 5B,C). The proportion of nascent to non-nascent UMIs was 18%, which echoes the proportion of unspliced to spliced UMIs identified using an equivalent E8.5 inDrop dataset^50^ (15%) (Extended Data Figure 5A).

**Figure 5.**
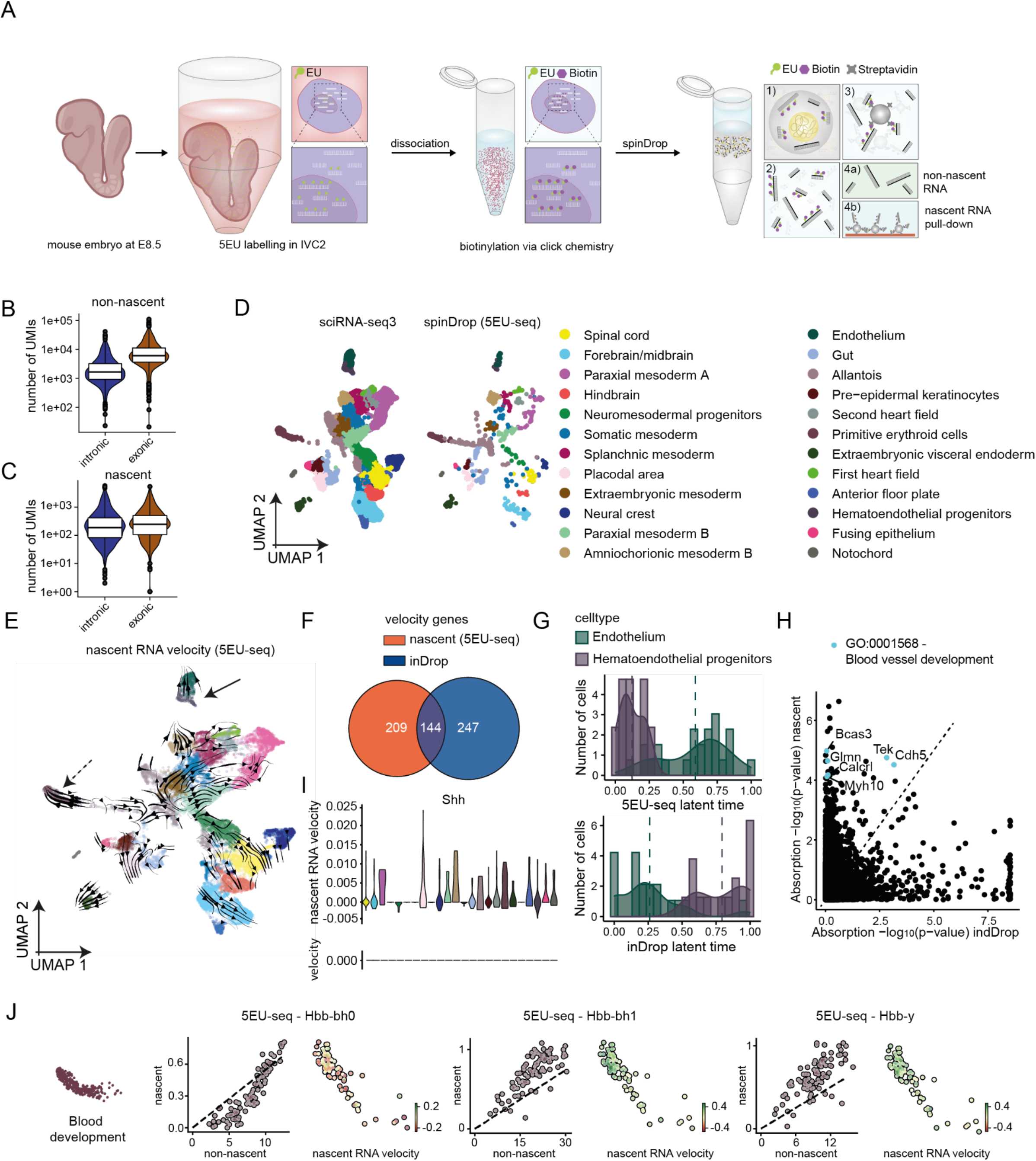
Nascent RNA sequencing using 5EU-seq defines transcription dynamics during mouse organogenesis A) Schematic of embryo processing at E8.5 using the droplet-based 5EU-seq method. The C57BL/6 embryos are incubated for 3 hours in IVC1 in presence of the 5EU analog. The cells are then dissociated, fixed, permeabilized and a biotin group is added using click chemistry. After processing with spinDrop, the recovered cDNA molecules are incubated with streptavidin-coated magnetic beads to separate nascent and non-nascent RNA/cDNA complexes. B) Number of UMIs detected for the non-nascent fraction mapping to either intronic or exonic regions. C) Number of UMIs detected for the nascent fraction mapping to either intronic or exonic regions. D) Integration of the spinDrop dataset with the sciRNA-seq3 mouse embryo dataset (downsampled to 20,000 cells) with the 5EU-seq data for label transfer shows capture of all main cell types. E) Single-cell nascent RNA velocity field using scVelo, projected on the shared embedding (spinDrop and sciRNA-seq3). The full arrow indicates endothelium maturation, the dashed arrow indicates blood maturation. F) Number of intersecting dynamical genes detected using scVelo in stochastic mode using unspliced and spliced matrices for the downsampled inDrop dataset (n=936 cells) and using the nascent and non-nascent RNA matrices with the spinDrop dataset. G) Latent time projections along the endothelial maturation trajectory, from haematoendothelial progenitors to endothelium, calculated using RNA velocity in the inDrop dataset or nascent RNA velocity in the spinDrop dataset. H) Lineage driver gene absorption probabilities for the endothelial maturation trajectory (-log10 transformed) calculated using Cellrank for the inDrop (RNA velocity) and spinDrop (nascent RNA velocity) datasets, highlighting an enrichment (p-value < 10^-5) for gene ontology terms relating to blood vessel development in the nascent RNA dataset. I) Nascent RNA and RNA velocity plots for *Shh* across the spinDrop and inDrop datasets. J) Nascent RNA sequencing uncovers novel haemoglobin transcriptional kinetics. From left to right, phase plots and velocity values superimposed on the UMAP projection for blood progenitors of *Hbb-bh0, Hbb-bh1* and *Hbb-y*.

Next, cell-type annotations from a sciRNA-seq3 dataset at E8.5 were projected on the spinDrop dataset as described in the previous section (Figure 5C). Because the spinDrop dataset was filtered to contain barcodes represented both in the nascent and non-nascent libraries, proportions of cell types compared to the sciRNA-seq3 reference may inform on the capabilities of the analog to diffuse throughout the embryo. For example, primitive erythroids, gut, endothelium and extra-embryonic mesoderm and endoderm cells were proportionally enriched in the 5EU-seq dataset, whereas neural cell types such as the spinal cord, forebrain/midbrain and hindbrain were depleted (Extended Data Figure 5D). Next, velocity vectors were computed via scVelo^51^ using the nascent and non-nascent transcripts as an input for the spinDrop dataset and unspliced and spliced transcripts for the equivalent inDrop dataset using and projected on a shared UMAP embedding (Figure 5E and Extended Data Figure 5E), yielding a list of dynamical genes with 144 genes intersecting and 456 non-intersecting genes (Figure 5F and Supplementary table 3). Major trajectories, such as the bi-potent commitment of neuromesodermal progenitors to spinal cord and paraxial mesoderm or the differentiation of haematoendothelial progenitors to endothelium could be observed with nascent RNA capture, whereas these trajectories were less clearly defined in the inDrop dataset (Extended Data Figure 5E). To quantify this finding, latent time values across the endothelial development trajectory were computed and accurately retraced natural development by computing velocity values using nascent RNA, whereas the trajectory was inverted in the inDrop dataset (Figure 5G) using conventional velocity measurements. In addition, the absorption probabilities computed using CellRank^52^ showed an enrichment for lineage drivers relating to blood vessel development (GO:0001568, FDR=3*10^-2) using nascent RNA sequencing (Figure 5H, Supplementary table 4), showcasing accurate delineation of developmental drivers. For some genes, RNA velocity did not capture expression dynamics altogether due to the lack of measurable unspliced molecules in the inDrop dataset. This can be explained by the stochastic nature of barcoded oligo(dT) probe binding to A/T-rich unspliced sequences across the transcript body^38^, hereby introducing a sequence-bias for velocity computation. In addition, the enrichment for 3’ molecules favours 3’-proximal intronic regions, which is highly gene-specific. Nascent RNA sequencing, on the other hand, performed better for the set of genes highlighted in Figure 5F because it does not rely on stochastic probe binding and bears no sequence bias, meaning that nascent transcripts are being read at the 3’ end of the transcripts similarly to non-nascent transcripts. One example is the morphogen *Shh* which was activated in the placodal area and notochord (Figure 5I), whereas RNA velocity could not be inferred in the inDrop dataset. Similarly, haemoglobin synthesis dynamics could exclusively be captured using nascent RNA sequencing, showing a repression of *Hbb-bh0* and induction of *Hbb-bh1* and *Hbb-y* in blood progenitors at E8.5 (Figure 5J). Some gene dynamics were detected both using nascent RNA sequencing and RNA velocity, but dynamics were not representative of natural development using the latter method, such as the erroneous split induction and repression of *Pax6* in the spinal cord using RNA velocity which was accurately identified as being solely inducted using nascent RNA measurements (Extended Data Figure 5F).

These findings position spinDrop as a modular methodology that may provide more comprehensive ‘-omics’ maps of heterogeneous tissues as current state-of-the-art methods.

## Discussion

spinDrop is an open-source droplet microfluidic workflow that uses droplet sorting and picoinjection to maximise *bona fide* information output from high-throughput single-cell experiments. We demonstrate that spinDrop enables deterministic sequencing of viable single-cells from the pool of empty droplets or droplets containing cell debris and damaged cells, which in turn reduces sequencing cost and removes artefacts from single-cell experiments. The method also supports cell type enrichment using fluorescently-labelled antibodies targeting cell surface markers, which may prove useful on samples where FACS cannot be employed due to cell number or viability concerns. In addition, gene detection rates were increased fivefold compared to inDrop^14^ to match the sensitivity of the leading commercial platform (10x Chromium). The cost bottleneck of high-throughput single-cell RNA-seq was addressed in spinDrop, with a 20 to 50% reduction in sequencing cost depending on input sample quality, and 6.2 decrease in library preparation cost compared to the 10x Chromium, leading to an average 60% decrease in overall cost per cell (0.4 USD per cell for spinDrop, 1USD per cell for 10x, Supplementary table 5)^12^. In addition, functionality across different input modalities was demonstrated for HEK293T cells, with high performance being achieved for whole cells, extracted nuclei and fixed cells. This will enable single-cell processing of archival tissues, which is a critical need in the clinic^41,53^. Although fixed cell processing has been demonstrated using the Drop-seq^54^ and 10x Chromium^55^, these applications suffer respectively from low cell capture (10-fold lower than spinDrop) and reliance on probe-based capture which is highly species specific and prevents genotyping applications which is crucial for investigating clinical samples. spinDrop was further demonstrated by profiling a damaged sample of the developing mouse brain at E10.5, removing the noise arising from empty droplets and damaged cells, thus showcasing the suitability of the method for molecular atlasing of complex and low input samples.

Application of quality control filtering methods to bioinformatically remove noise from single-cell datasets is standard procedure before performing downstream analysis^56^. However, several drawbacks are associated with computational filtering of noisy cells from datasets. For example, damaged cells and cell debris can be filtered out using gene expression counts mapping to mitochondrial RNA molecules, as they are retained at higher rates than cytoplasmic RNA when the cell membrane is perforated. However, mitochondrial RNA content varies with cell type and species^57^, which means that filtering on pre-set thresholds may alter the dataset and bias subsequent interpretation. Another metric to identify empty droplets and damaged cells in a dataset is generated by computing the nuclear fraction ratio of unspliced-to-spliced reads^19^. Because empty droplets contain higher levels of spliced cytoplasmic RNA and damaged cells contain higher levels of unspliced nuclear RNA, the latter can be filtered out by setting thresholds on nuclear fraction (ratio of unspliced to spliced UMI counts). However, nuclear fractions may, again, differ widely between the profiled cell types. For example, erythrocytes contain small traces of mature RNA molecules and may therefore be identified as empty droplets using current tools^19^. Fluorescence-activated sorting of droplets containing viable cells at the moment of encapsulation circumvents this issue and deterministically resolves viable cells, independently of mitochondrial RNA content or cell type. Another capital pre-processing step consists in the removal of cellular doublets from the dataset. However, current tools have variable performance depending on cell-type and mostly identify heterotypic doublets^24,25^ (i.e. multiplets from different cell types). Here, we demonstrate proof-of-principle for discarding doublets using FADS by applying an upper threshold on fluorescence. Sorting metrics showed reduced co-encapsulation rates compared to predictions, illustrating the potential of the method to remove doublets from the sequenced pool. Deterministic sequencing of droplets containing single-viable cells or intact nuclei therefore reduces artefacts introduced by downstream processing, but also alleviates the sequencing cost associated with these artefacts.

Although some methodologies have utilised droplet sorting for single-cell sequencing applications, they are limited in their applicability to molecular atlasing. Studies by Clark *et al*.^*58*^ and Zhang *et al*.^*59*^ are primarily utilising sorting for cell-type specific isolation rather than the enrichment of single viable cells^16^. Machine vision methods have been described for the deterministic sorting of co-encapsulated single-cells and single barcoded microgels, but current workflows suffer from low throughput (77-fold smaller than spinDrop)^60^ and do not take cell viability into account during sorting^60^. Furthermore, none of the aforementioned methodologies have been designed to increase sensitivity compared to other methods to match the leading commercial standard (10x Chromium), and the protocols have not been demonstrated across input modalities for the processing of archival tissues and nuclei, thus limiting their applicability towards molecular atlasing endeavours across sample types. In addition, the advantages of excluding damaged cells and empty droplets were not explored using these methodologies.

More generally, we anticipate that the powerful combination of droplet sorting and picoinjection described here will complement microfluidic methods that go beyond transcriptome sequencing in droplets, such as ATAC-seq^32^ or multi-omic profiling^61^. We demonstrate the benefit of multi-step microfluidics by performing reverse-crosslinking in droplets, which allowed us to profile nascent RNA transcription during mouse organogenesis using 5EU-seq implemented in droplets, uncovering previously hidden layers of biology that are not attainable using state-of-the-art methods. Integration with commercial toolboxes like the 10x Chromium will present a unique opportunity to decrease cost and increase customer acquisition further. Future droplet sorting implementations may include multimodal sorting, such as fluorescence coupled to image-based sorting^62^ to further decrease doublet rates and acquire cell-type phenomic profiling, which would enable higher cell loading concentrations and increased throughputs as well as multimodal phenotypic characterization.

Overall, spinDrop is well aligned with atlasing efforts like the Human Cell Atlas, as it provides ‘ground truths’ and sensitive reference datasets of human biology. In addition, the method is the first to unify low-cost, high-throughput and high-sensitivity, which is critical to the success of the community’s quest for accurately determining the spectrum of heterogeneity in cell biological scenarios.

## Methods

### Design of the droplet generation device

The integrated device for compressible barcoded bead and single-cell co-encapsulation droplet generation device followed FADS (Extended Data Figure 1A) is a modified droplet generator used in previous studies^14,28,37^ with an added fluorescence-activated droplet sorter (FADS) for enrichment of droplets containing single viable cells stained with Calcein-AM. Our FADS module is based on integrated fibres^63^ for both excitation and detection of fluorescence. The emission light emerges from the detection optical fibre connected to the detector tube housing a set of emission filters mounted before the detector of the photomultiplier tube (Extended Data Figure 2A). When a fluorescence light signal was higher than an arbitrarily set threshold, a high voltage pulse was generated (1 kV) and delivered to the microfluidic sorting junction by ‘salt electrodes’ filled with 5M NaCl solution. As a result, highly fluorescent droplets with live cells were derailed to the collection channel for positive ‘hits’. The cell encapsulation and sorting chip comprise sections of increasing depths: i) the droplet generator is 75 µm deep to ensure good encapsulation efficiency for 60 µm barcoded beads, ii) the depth of the detection spot (95 µm) is determined by the width of the fibres, and iii) the sorting junction was made deeper (175 µm) in order to avoid squeezing of droplets and facilitate their redirection into the negative channel. The microfluidic device additionally comprised two inlets (number 6 and 7) and a flow-focusing junction for generation of ‘buffer droplets’ that did not contain cells, beads or ambient RNA. The role of buffer droplets was to facilitate the handling of droplets for the subsequent picoinjection step and increase the volume of the aqueous phase after sorting for easier handling.

The picoinjection module is an improved version of a device used in our previous study^37^. The inlet for emulsion diluting oil (number 2, Extended Data Figure 1B) is located this time behind the droplet emulsion inlet (number 1, Extended Data Figure 1B) which reduces the fragmentation of droplets in densely packed emulsion before reinjection. The final section of the picoinjection device was made deeper (180 µm) to stabilise droplets and reduce their merging during sudden change of depth between the shallow microfluidic channel and a wide collection tubing.

### Photolithography of microfluidic moulds

The channel layout for the microfluidic chips was designed using AutoCAD (Autodesk) and printed out on a high-resolution film photomask (Micro Lithography Services). The designs in Extended Data Figure 1 are deposited on https://openwetware.org/wiki/DropBase:Devices and can be found in the supplementary file ‘SI_spinDrop CAD designs_5 masks’. The microfluidic devices were fabricated following standard hard lithography protocols that can be performed in local cleanrooms or outsourced to contract manufacturing companies. First, microfluidic molds were patterned on 3” silicon wafers (Microchemicals) using high-resolution film masks (Microlithography Services Ltd) and SU-8 2015, 2075 and 2100 photoresists (Kayaku Advanced Materials). A MJB4 mask aligner (SÜSS MicroTec) was used to UV expose all the SU-8 spin-coated wafers. The thickness of the structures (corresponding to the depth of channels in the final microfluidic devices) was measured using a DektakXT Stylus profilometer (Bruker).

### Microfluidic PDMS Chip Fabrication

The settings used for photolithography can be found in Supplementary table 6 and 7.

### Soft lithography

To manufacture PDMS microfluidic devices, 20-30 grams of silicone elastomer base and curing agent (Sylgard 184, Dow Corning) were mixed at a 10:1 (w/w) ratio in a plastic cup and degassed in a vacuum chamber for 30 minutes. PDMS was then poured on a master wafer with SU-8 structures and cured in the oven at 65 °C for at least 4 hours. Next, the inlet holes were punched using two types of biopsy punchers with plungers (Kai Medical): a 1.5 mm diameter punch was used to make the inlet for cell delivery tip, number 2 (Extended Data Figure 1A), outlet tip for droplet collection, number 9 (Extended Data Figure 1A) and the inlets for droplet reinjection, number 1 (Extended Data Figure 1B), while other inlets were made using a 1-mm-wide biopsy puncher. The FADS PDMS chip was then plasma bonded first to an approximately 1-mm thin PDMS slab (cured beforehand) and then to a 52 mm x 76 mm x 1 mm (length x width x thickness) glass slide (VWR) in a low-pressure oxygen plasma generator (Femto, Diener Electronics). As a result, we obtained a 3-layer device with a patterned PDMS on top, a thin PDMS slab in the middle, and a glass slide at the bottom. The picoinjection chip was bonded only to the glass slide and the resulting device had two layers. Next, the hydrophobic modification of microfluidic channels was performed by flushing both types of devices with 1% (v/v) trichloro(1H,1H,2H,2H-perfluorooctyl)silane (Sigma-Aldrich) in HFE-7500 (3M) and baked on a hot plate at 75 °C for at least 30 minutes to evaporate the fluorocarbon oil and silane mix. Next, the fabricated FADS device was mounted on the microscope stage for the assembly of fibres and tubing and the picoinjection chip was used in the second stage of the microfluidic experiment.

### Assembly of the fibre FADS chip

The fabrication of the sorting chip required additional integration with multimode optical fibres for delivering excitation light and detection of fluorescence signals. Consecutive steps of the assembly of a FADS chip are presented in Extended Data Figure 2A. The excitation light fibre (cat. no M94L02, Thorlabs) had a cladding diameter of 125 µm and a core diameter of 105 µm with a numerical aperture (NA) of 0.1. The detection fibre (cat. no M43L02, Thorlabs) had a cladding diameter of 125 µm and a core diameter of 105 µm with NA of 0.22. Both fibres were cut at their ends to remove one of the FC/PC connectors, and next, the outer protective PVC jacket was removed using a three-hole fibre stripper (cat. no FTS4, Thorlabs). The Kevlar protective threads were cut with a scalpel and finally acrylate coating was removed using a fibre stripping tool (cat. no T06S13, Thorlabs). In the next step, the tip of the fibre tip was cleaved using a ceramic fibre scribe (cat. no CSW12-5, Thorlabs) in order to obtain a flat tip end. The quality of the cleavage was inspected by passing a low power (e.g. mW) laser light through the fibre and a visual inspection of the shape of a beam emerging from the fibre tip end. If necessary, the cleavage was repeated until the spherical shape of the light beam was observed. Fixing of fibres to the chip was performed on the microscope stage and a microscope camera was used to verify the position of the fibre ends. First, the microfluidic channels housing the fibres were filled with HFE-7500 oil and then fibre tips were manually inserted into the chip. fibres were stabilized by attaching them to the glass slide with pieces of insulation tape.

### mESC and HEK293T cell culture and preparation

HEK293Ts were passaged every second day and cultured in T75 flasks. The culture media was DMEM (4500 mg/L gluc & L-glut & Na bicarb, w/o Na pyr, D5796-500ML, Sigma) supplemented with 10% heat-inactivated FBS and 1x Penicillin-Streptomycin. For passaging and collection, the cells were washed with 10 mL ice-cold 1x PBS (Lonza) twice. 9 mL of PBS was added to the flask and cells were detached by adding 1 ml of 10x Trypsin-EDTA (Sigma-Aldrich) and incubated at 37 °C for 5 minutes. Trypsin-EDTA was then inactivated with 15 mL of DMEM 10% FBS and cells were incubated at 37 °C for 5 minutes. The cells were then pelleted at 300g for 3 minutes and the supernatant was aspirated. For the experiment, 1 µL of Calcein-AM and 1 µL of ethidium homodimer-1 were added to one millilitre of washed HEK293T cells and incubated on ice for 25 minutes. The cells were then pelleted at 500g for 5 minutes at 4 °C and resuspended in 1x PBS, and brought to a concentration of 250 cells per µL for 1x loading and 1,250 cells per µL for 5x loading. The cells were then mixed 1:1 with a solution of 1x PBS + 30% (v/v) Optiprep for encapsulation.

Wild type mouse embryonic stem cells were cultured in 2i+LIF medium (DMEM/F-12 without L-glutamine (Gibco) and Neurobasal medium without L-glutamine (Gibco) in a 1:1 ratio, supplemented with 0.1% sodium bicarbonate (Gibco), 0.11% Bovine Albumin Fraction V Solution (Gibco), 0.5x B27 supplement (Gibco), 1x N2.BV (Cambridge Stem Cell Institute), 50 µM 2-mercaptoethanol (Gibco), 2 mM L-glutamine (Gibco), 1x Penicillin-Streptomycin (Gibco), 12.5 µg/mL human insulin recombinant zinc (Gibco), 20 ng/mL leukaemia inhibitor factor (Cambridge Stem Cell Institute), 3 µM CHIR99021 (Cambridge Stem Cell Institute), 1 µM PD0325901 (Cambridge Stem Cell Institute)) at a cell density of ∼8,000 cells/cm^2^. To passage the cells or generate a single cell suspension, the cells were treated with Accutase for 3 minutes at 37 °C and subsequently washed with 10x the volume wash medium (DMEM-F12 + 1% BSA), centrifuged at 300g for 3 minutes and then resuspended in either culture medium or 1x PBS. For experimental procedures the cells were then processed similarly to the HEK293T cells.

### Preparation of mESC and HEK293T nuclei

The cells were cultured and harvested as described in the previous paragraph. The suspension of nuclei was obtained following the Nuclei EZ prep guidelines applied to cells resuspended in PBS and stained for 20 minutes on ice by supplementing 1 µL of Vybrant DyeCycle Green DNA stain to the nuclei resuspended in 1x PBS, 7.5% Optiprep and 0.04% BSA.

### mESC and HEK293T nuclei or whole cell species mixing library preparation

The cell solution was loaded at 125 HEK293T cells/μL and 125 mESCs cells/μL or 125 HEK293T nuclei/μL and 125 mESCs nuclei/μL. Subsequent droplet generation and sorting was conducted as described above.

### Preparation of cell suspensions from the developing mouse brain at E10.5

#### a. Ethics statement

Mice maintenance and care was performed according to national and international guidelines. All experiments were in accordance with the regulations of the Animals (Scientific Procedures) Act 1986 Amendment Regulations 2012 and were approved by the University of Cambridge Animal Welfare and Ethical Review Body (AWERB) and by the Home Office.

#### b. Animal maintenance and neural embryonic tissue retrieval

We inspected animals daily and those with any sign of health concerns were culled immediately by cervical dislocation. All experimental mice were free of pathogens and were housed on a 12-12 hour light-dark cycle and they had unlimited access to water and food. Temperature in the facility was controlled and maintained at 21°C. Mice for post-implantation embryo recovery (CD-1 females and males from Charles River were acclimated for 1 week prior to use) were utilised from 6 weeks of age. Females and males were mated for up to five days or until a plug was found; inspection for vaginal plugs was performed daily. Females were culled by cervical dislocation at the appropriate time point to obtain embryos of the correct age. The day on which a vaginal plug was found was scored as E0.5. Embryo dissection was carried out in M2 medium (Sigma). Mouse embryos were carefully collected by cutting open the decidua and the yolk sac. To collect the neural tissue, the pial meninges were removed and the head was cut above the eyes to collect the forebrain, midbrain and part of the hindbrain.

#### c. sample dissociation

The collected tissue was quickly washed in PBS twice and centrifuged for 5 minutes at 0.2 x g prior to dissociation with 500 μL of TrypLE Express (Gibco) at 37 °C. The tissue was allowed to dissociate for 15 minutes, with pipetting for 10-15 times (avoiding bubble formation) every 5 minutes to help tissue breakdown. If clumps of cells persisted, dissociation was continued for an additional 5 minutes. Dissociation was halted by adding 2 mL of basal DMEM medium supplemented with 15% of foetal bovine serum. The cell suspension was filtered through a 40 um cell strainer (Greiner), centrifuged, washed once with 1x PBS, centrifuged again and resuspended again in a small volume of PBST (PBS with 0.02% Tween-20) and stored on ice until encapsulation. After the PBS wash, a small aliquot of the cell suspension was used to assess viability by mixing it in a 1:1 ratio with Trypan Blue (Sigma) and quantified using a haemocytometer.

#### General description of the microfluidic encapsulation, sorting and picoinjection

##### Co-encapsulation of cells and barcoded beads followed by droplet sorting

A detailed protocol^28^ was used as a reference for droplet generation. Here we present step-by-step guidelines for performing cell encapsulation and droplet sorting:

1. First, the microfluidic fibre FADS chip was installed and on the stage of an inverted microscope (Olympus XI73), and the optical fibres were inserted as presented in Extended Data Figure 2A, numbers 1-3.
2. Next, syringes containing 5 M NaCl were connected to the device (Extended Data Figure 2A, number 4) and the electrode channels pre-filled with salt solution filtered as previously described^31^.
3. Three 2.5-mL or 5-mL glass syringes (SGE) were filled with 5% (w/w) 008-FluoroSurfactant (RAN Biotechnologies) in HFE-7500 (3M) and connected to oil inlets (Extended Data Figure 2A, number 5).
4. Next, three pieces of polyethylene tubing (Portex, Smiths Medical) were connected to two 1-mL gas-tight syringes (SGE or Hamilton) and filled with PBS (Lonza). The tubing was manually filled with PBS, and a small, 1-cm-long air bubble was left at the end tip of each tubing. The aqueous solution for buffer droplets was 1x First Strand buffer (Invitrogen), 4.2 mM DTT (Invitrogen). The lysis mix and buffer droplets mix were manually aspirated into the tubing, and the small air bubble provided a separation between the reagents and the PBS buffer.
5. Then, 150 μL of cell suspension was manually aspirated into the cell loading tip pre-filled with mineral oil (Sigma-Aldrich). Detailed fabrication protocol of cell delivery and droplet collection tips is provided in our recent article^37^.
6. Next, all three tubings and the cell chamber with cell suspension were inserted to the corresponding inlets of the droplet generation chip (Extended Data Figure 2A, number 6).
7. Droplet collection tip^37^ (Extended Data Figure 1C) and long outlet tubing were connected to the positive and negative outlets, respectively (Extended Data Figure 2A, number 7).
8. Finally, in dark conditions, the barcoded beads were aspirated into the tubing which was next connected to the microfluidic device (Extended Data Figure 2A, number 8). The inDrop barcoded beads were prepared prior to the experiment according to the inDrop protocol^28^, with the inDrop v3 oligonucleotide barcoding scheme^29^. Just before aspiration, the beads were washed three times in 10 mM Tris-HCl (pH 8), 0.1 mM EDTA and 0.1% Tween-20 and resuspended in 55 mM Tris-HCl (pH 8), 0.1% (v/v) IGEPAL CA-630, 75 mM KCl, 0.05 mM EDTA (Ambion), 0.05% Tween-20. The lysis mix was as follows: 120 mM Tris-HCl (pH 8), 3.15 mM dNTPs (each), 0.6% (v/v) IGEPAL CA-630, 7.2 U/mL Thermolabile proteinase K (NEB).
9. The solutions were injected in the in-line microfluidic device using neMesys pumps (Cetoni) and sorted with the following flow rates: 150 µL/hr for the lysis mix, 150 µL/hr for the cell mix, 60 µL/hr for the beads, 700 µL/hr for the main oil, 2,500 µL/hr for the spacing oil to generate droplets with average volume 1.3 nL with around 75 Hz frequency. Additional flows of 15 µL/hr for aqueous phase and 30 µl/hr for oil were applied to generate buffer droplets.
10. Next, the fluorescence was recorded in high throughput and droplets were sorted according to the set threshold. After stabilisation of a droplet flow, the syringe with mineral oil was removed from the collection tip outlet tubing and the droplet sorting began. A short 1-ms-long pulse was delivered with a delay of 3.5 ms to the microfluidic device by ‘salt electrodes’ filled with 5M NaCl solution and, as a result, highly fluorescent droplets with single cells were derailed to the collection channel for positive ‘hits’ (b). The duration and delay of pulse can be modified according to the flow rates and the desired throughput of the sorting (usually around 75 Hz), depending on the droplet generation rate. Sorting device was used only once for each experiment since the droplet generation was performed on the same chip.
11. The positive and buffer droplets were collected in a collection chamber pre-filled with mineral oil and incubated at room temperature (23 °C) for 25 minutes. The barcodes were then solubilized from the bead via UV exposure using a high-intensity UV inspection lamp (UVP) for 7 minutes.
12. The collection chamber was then immersed in a water bath solution at 70 °C for 10 minutes and directly immersed in an ice-cold recipient (half part water and half part ice) for 5 minutes after the incubation was finished.

#### Picoinjection of RT mix

Before starting the picoinjection of droplets containing single-cell lysates, the electrode sections (Extended Data Figure 1B) of the devices, were pre-filled with filtered 5 M NaCl as previously described. The picoinjection chip was filled with 5% (w/w) 008-FluoroSurfactant (RAN Biotechnologies) in HFE-7500 (3M) using a pre-filled 2.5-ml glass syringe (SGE) connected to a piece of tubing (Portex, Smiths Medical). The reaction mix was primed, and the tip containing the emulsions (with fluorinated oil evacuated by pushing the glass syringe until the emulsions reached the exit of the tip) was primed and connected to the device. The droplets were then re-injected in a pico-injector device and coalesced with a RT solution at 1:1 ratio of flow rates. The RT solution was 1.8x First Strand Buffer (5x FS, Invitrogen), 2.52 mM MgCl2 (Ambion), 9 mM DTT (Invitrogen), 3.6 U/µL RnaseOUT (Invitrogen), 24 U/µL SuperScript III (Invitrogen). For testing the effect of molecular crowding on RNA capture, 13.5% PEG8000(NEB) was added to the RT solution. For RT conditions using the Maxima H-, the following mixture was used: 1.8x RT buffer (Invitrogen), 2.5 mM MgCl2, 2 U/µL RnaseOUT (Invitrogen) and 18 U/µL Maxima H-RT (Invitrogen) and the RT incubation temperature was switched to 42 °C. The flow rates were the following: 200 µL/hr for the droplet mix, 200 µL/hr for the RT solution, 40 µL/hr for the spacing oil and 400 µL/hr for the main oil. The libraries were then prepared as per the inDrop library preparation protocol and sequenced on a Nextseq 75 bp High Output Illumina kit (Read 1: 61 cycles, Index 1&2: 8 cycles each and Read 2: 14 cycles).

#### Processing of mouse C57BL/6 PBMCs for cell-type enrichment

Frozen Splenocyte vials from C57BL/6 mouse were thawed in a water bath at 37 °C and immediately pipetted in 13 ml of pre-warmed (37 °C) 1xPBS, 10% FBS. The mix was spun down at 300g for 5 minutes at 4 °C and washed twice with 1x PBS 2% FBS. The cells were then strained through a 50 µm cell strainer and 1 million cells in 100 µL were incubated with 10 µL of Fc receptor block (Miltenyi Biotec) for 10 minutes on ice. 2 µL of PE-anti IgM, CD19 and CD45R antibodies (Miltenyi Biotec) were added to the mixture and the cells were further incubated on ice for ten minutes. The cells were then washed two additional times with ice-cold PBS and counted and encapsulated using the native inDrop conditions. The light source used for FADS-sorting of B-cells was a diode-pumped solid-state (DPSS) OBIS 561 nm 50 mW laser (Coherent), and the emission was captured through one notch filter 561 nm and two bandpass 593/40 nm filters. The droplets from each sorting channel were imaged under an EVOS FL fluorescence microscope. The libraries were then processed as described in the previous paragraph.

#### Processing of mouse embryos at E8.5 for nascent RNA sequencing using 5EU-seq

The embryos were recovered as described in the previous sections and placed in IVC1^64^ media for 3 hours in presence of the 500 µM 5EU (5-ethynyl-uridine). The embryos were cut in smaller pieces using sharp blades, and were dissociated with TrypLE for 5 minutes at 37°C. The reaction was quenched using DMEM/F-12 supplemented with 10% FBS and the cells were fixed and pre-processed as per the scEU-seq protocol^49^. The cells were then resuspended in PBS, counted and processed through the spinDrop workflow without fluorescence-activated droplet sorting. The downstream library preparation protocol after de-emulsification follows the methodology described in the scEU-seq methodology, however, the final PCR amplification was performed as per the spinDrop protocol, to account for differences in sequencing adapters.

#### Computational analysis

The bcl files were converted to fastq using bcl2fastq (Illumina) and quality controlled using FastQC^65^ and demultiplexed using Pheniqs^66^. For benchmarking of HEK293T cells, the biological read from the 10x v2 dataset was trimmed to 61 base pairs to match the length of the reads for the libraries prepared with spinDrop. Both mouse (GRCm38, ensembl 99 annotations) and human (GRCh38, ensembl 99 annotations) genomes were indexed using STAR^67^ and gene expression matrices were generated using zUMIs^43^. Gene names and biotypes were queried from bioMart^68^ and downstream integration of datasets and tertiary analysis were performed using the Seurat v3^45^ package. For downsampling measurements and intronic and exonic repartition calculations, the matrices for each coverage were obtained from the dgecounts rds object generated from zUMIs. Quality control on cell barcodes was achieved using DropletQC v1.0^19^ on unfiltered matrices generated using zUMIs. For integration of the 10x v1^44^, spinDrop and sciRNA-seq3^11^ mouse brain datasets, the *FindTransferAnchors* function from Seurat v3 was used to create a shared embedding. To transfer the annotations from the 10x v1 to the other datasets, the Louvain algorithm was used (*FindClusters* function) to define clusters in the shared embedding. The label was then transferred to the shared embedding using a maximum Pearson correlation coefficient calculation between the average expression of the 10x v1 dataset labels and the corresponding cluster ID from the shared embeddings. Differential gene expression analysis was then conducted using a Wilcoxon rank sum test (*FindMarkers* function).

#### Data availability

The 1:1 3T3 and HEK293T mixture 10x Chromium v3 dataset used for benchmarking HEK293T cells is available on their website in the ‘Datasets’ category. The sciRNA-seq3 E10 dataset was obtained from http://tome.gs.washington.edu/, and the 10x v1 mouse brain dataset was downloaded from SRA with accession number PRJNA637987.

The microfluidic chip designs in Extended Data Figure 1A can be found in our repository DropBase (https://openwetware.org/wiki/DropBase:Devices)

#### Code availability

Code is available at https://github.com/droplet-lab/spinDrop

## Acknowledgments

J.D.J. received scholarship support from the BBSRC, T.S.K. held an EU H2020 Marie Skłodowska-Curie Individual Fellowship (MSCA-IF 750772), A.L.E. was supported by the Cambridge Trusts and the EU H2020 Marie Curie ITN MMBio and T.N.K. by an AstraZeneca studentship. M.T. was supported by the International Centre for Translational Eye Research (project MAB/2019/12, carried out within the International Research Agendas programme of the Foundation for Polish Science co-financed by the European Union under the European Regional Development Fund). This work was supported by the EU Horizon 2020 programme (ERC Advanced Investigator Awards to F.H., 69566 and M.Z.G., 669198), the Wellcome Trust (WT108438/C/15/Z to F.H. and 207415/Z/17/Z to M.Z.G.) and the NIH (Pioneer Award to M.Z.G., DP1 HD104575-01). The authors would like to thank the members of the Hollfelder laboratory for their feedback. We thank Dr. Anna Alemany for help and suggestions for the data analysis.

## Author information

J.D.J., T.S.K. and F.H. conceptualised the study. T.S.K. and J.D.J. developed and optimised the droplet microfluidic workflow. J.D.J. developed and optimised the molecular workflow. J.D.J., A.L.E. and T.N.K. retrieved the cultured cells. G.A., C.H. and J.D.J. retrieved and processed the mouse embryos. J.D.J., T.S.K. and D.B.M. performed the encapsulations. J.D.J. performed library preparation and sequencing.

J.D.J. and M.T. performed downstream analysis of sequencing results. J.D.J., T.S.K. and F.H. wrote the manuscript, with input from all authors. F.H., G.M.F., S.T. and M.Z.G. supervised the work.

## Supplementary data

**Supplementary table 1** Wilcoxon rank sum two-sided differential expression test between 10x and spinDrop using HEK293T cells. Bonferroni-adjusted p-values and uncorrected p-values are provided.

**Supplementary table 2** Wilcoxon rank sum two-sided differential expression test between 10x and spinDrop using neuroblast cells from the mouse brain atlas. Bonferroni-adjusted p-values and uncorrected p-values are provided.

**Supplementary table 3** List of dynamical genes in the 5EU-seq and inDrop datasets computed using scVelo.

**Supplementary table 4** Fisher two-sided test computing the lineage drivers for endothelium specification for the 5EU-seq and inDrop datasets.

**Supplementary table 5** Cost per cell calculation to run a spinDrop experiment for the equipment, sequencing and library preparation.

**Supplementary table 6** Photolithography protocol details for the FADS microfluidic device.

**Supplementary table 7** Photolithography protocol details for the picoinjector microfluidic device.

**Supplementary video 1** FADS sorting in the positive channel of a droplet containing a single viable cell.

**Supplementary video 2** Picoinjection of a reverse transcriptase mix into the incoming droplets.

**Extended Data Figure 1.**
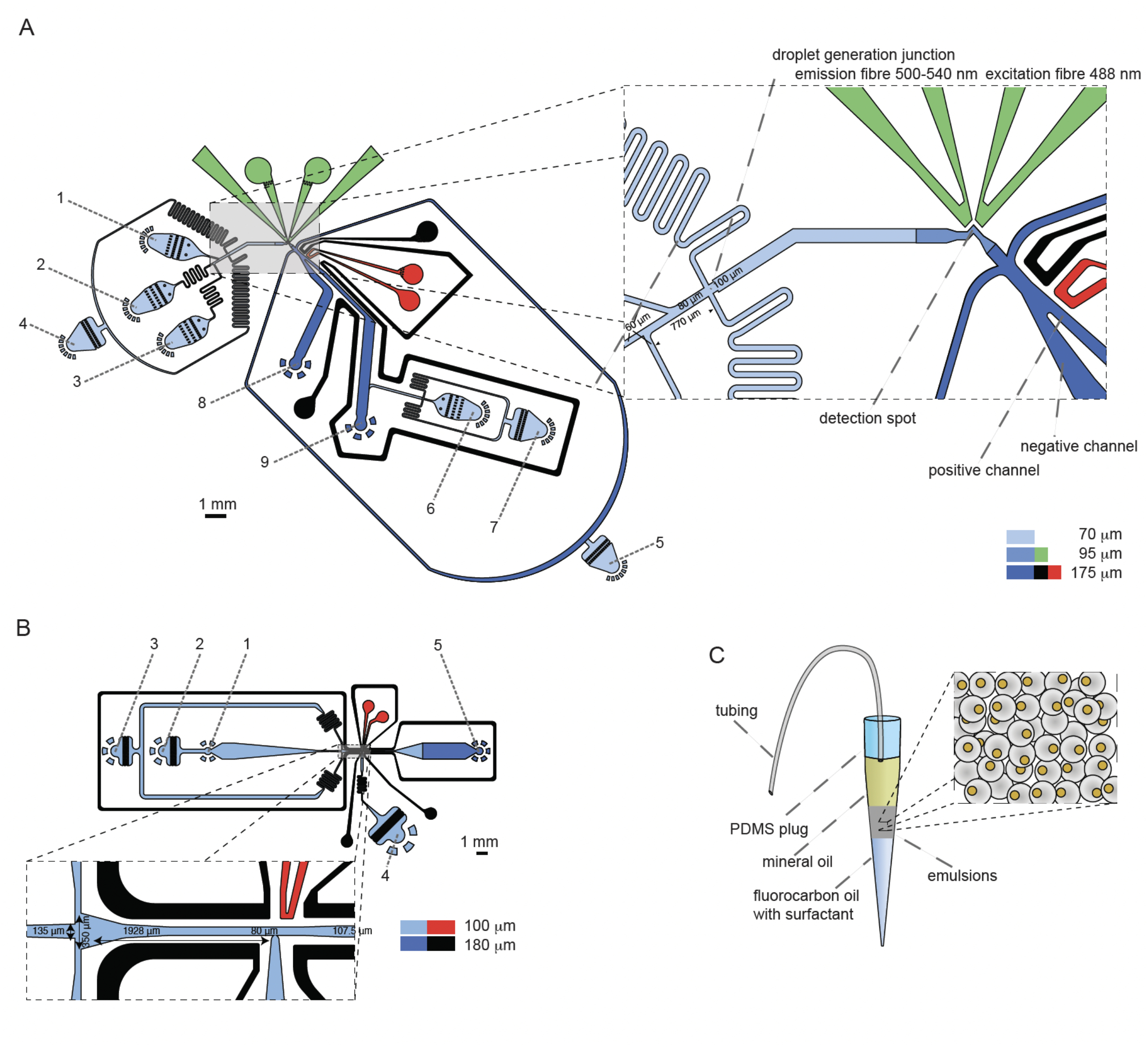
Schematics of the microfluidic devices **A)** Blue colour denotes fluidic channels, black – channels for ground electrode, red – signal electrode and the green colour indicates auxiliary channels for insertion of optical fibres. Design of the integrated device used for barcoded bead and single-cell co-encapsulation followed by fluorescence-activated droplet sorting. 1) Input channel for the barcoded compressible microgel loading. 2) Input channel for the cell loading, 3) input channel for, the lysis mix, 4, 5, 7) input channel for the fluorinated oil with admixed surfactant, 6) input channel for aqueous solution for generation of buffer droplets. 8) Outlet for negative droplets, 9) outlet for positive droplets. **B)** Design of the picoinjector device, used to inject the RT enzyme mix. 1) Input channel for droplet reinjection, 2) emulsion diluting oil inlet, 3) droplet spacing oil inlet, 3) inlet for the RT mix to be picoinjected, 5) droplet exit channel. C) Scheme of the container tip used for droplet collection and reinjection in the picoinjector device. The tip is connected to a glass syringe which enables aspiration and delivery of emulsions. The tip can be connected to a PDMS plug to close the system for incubation in the water bath.

**Extended Data Figure 2.**
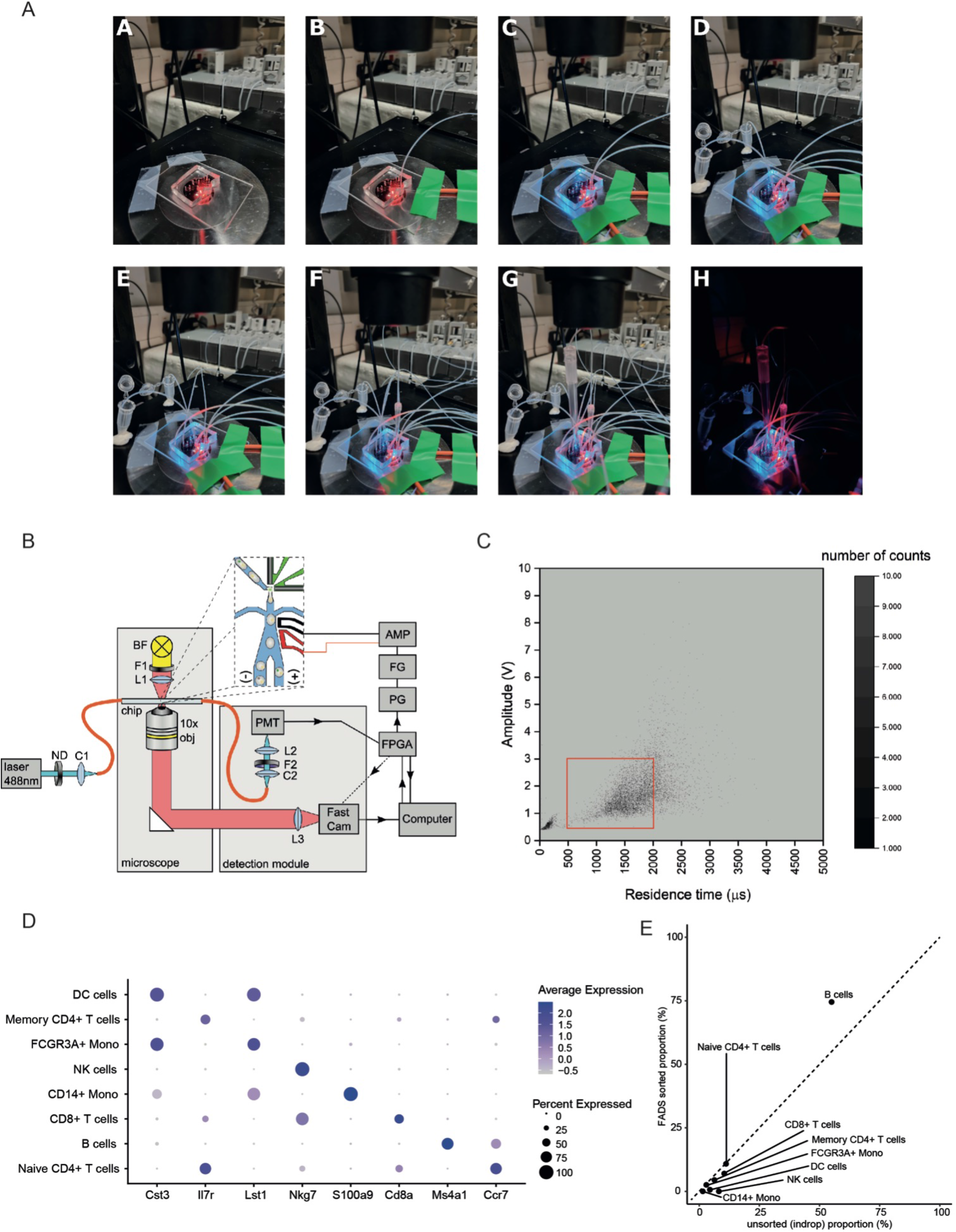
FADS set-up and sorting Photographs presenting consecutive steps of FADS chip assembly. A) Mounting of the chip on the microscope stage. B) Insertion of the detection fibre. C) Insertion of the excitation fibre. D) Connecting tubing with NaCl solution and filling of salt electrodes. Please note that outlet tubings do not touch the microscope, and they are placed inside Eppendorf tubes E) Connecting tubing with lysis mix, buffer and oil/surfactant mixtures. F) Insertion of cell loading tip. G) Insertion of droplet collection tip and the negative outlet tubing. H) Insertion of the tubing with barcoded beads. Final device ready for encapsulation in a dark room required to protect UV-cleavable beads. Refers to Extended Data Figure 1A with design. **B)** Scheme of the optical setup used for fluorescent activated droplet sorting for cells stained with Calcein-AM. The setup is a modification of a previously presented system for fluorescence-activated droplet sorting (10). The large part of the microfluidic device was illuminated by the red light coming from the microscope lamp through a 593nm longpass filter F1 (FF01-593/LP-25, Semrock) and a lens of the microscope condenser (L1). Next, the transmitted red light was collected by the objective (usually 10x or 20x), leaving the microscope through the camera port to be recorded by a fast camera (Miro eX4, Phantom, USA). The light from the 488 nm laser (Vortran Stradus) passed through a neutral density (ND) filter OD=0.5 (NE05A,Thorlabs). Next, the light was delivered to the sorting junction by coupling the laser beam via C1 collimator (adjustable FC/PC collimator, CFC-11X-A, Thorlabs) to the incident light optical fibre (105 µm, 0.1 NA, FC/PC, Thorlabs). Usually, the input intensity of the 488nm laser was set to 10 mW, which translates to approximately 3 mW on the chip, after passing the ND filter. The light emerging from the detection optical fibre (105 µm, 0.22 NA, FC/PC Thorlabs) was connected via a C2 collimator (CFC-11X-A, Thorlabs) to the detector tube housing the F2 set composed of 1-notch (NF488-15, Thorlabs) and two bandpass filters (FF01-550/88-25, Semrock) mounted 50 mm before the detector of photomultiplier tube PMT (PM002, Thorlabs). Simple spherical lenses (L2, L3) were used to focus the light on camera’s detectors and PMT. The fluorescent light signal was recorded by the PMT coupled to a FPGA NI card and analysed by a custom written LabVIEW program in real-time. For sorting of cells labelled with IgM-PE antibodies a different laser (Coherent OBIS 561 nm, 50 mW) and other filters were used: F1 - 635 nm longpass filter (BLP01-635R-25, Semrock), F2 - a set composed of one longpass filter 561 nm (BLP02-561R-25, Semrock) and two bandpass 593/40 filter (FF01-593/40-25, Semrock). The input intensity of the 561 nm laser was set to 25 mW, resulting in approximately 8 mW on the chip, after passing the ND filter. Black arrows indicate the direction of signals and triggers in the system. When a fluorescent light signal was higher than an arbitrarily set threshold, the FPGA card triggered the generation of a high voltage pulse (1 kV) by the series of electronic devices: pulse generator PG (TGP110, Thurlby Thandar Instruments) to function generator FG (TG200, Thurlby Thandar Instruments) to high voltage amplifier AMP (610E, Trek). **C)** FADS sorting dataset showing the recorded fluorescent signal data points collected across an experiment. The sorted droplets in the red rectangle (intensity 0.5-3V, time 500-2000 us) represent viable HEK293T cells stained with Calcein-AM, and excludes the background empty droplets (bottom left) and droplets containing cell doublets with high fluorescence signal (right handside). **D)** DotPlot for cell type-specific marker genes for the shared embedding between the 10x v2, inDrop FADS sorted for B-cells (using CD45R, CD19 and IgM-PE labelled antibodies), and the inDrop datasets generated from frozen C57BL/6 PBMCs cells. **E)** Cell type proportional representation between the inDrop FADS sorted B-cells and unsorted inDrop PBMCs dataset showing an enrichment for B-cells when using FADS.

**Extended Data Figure 3.**
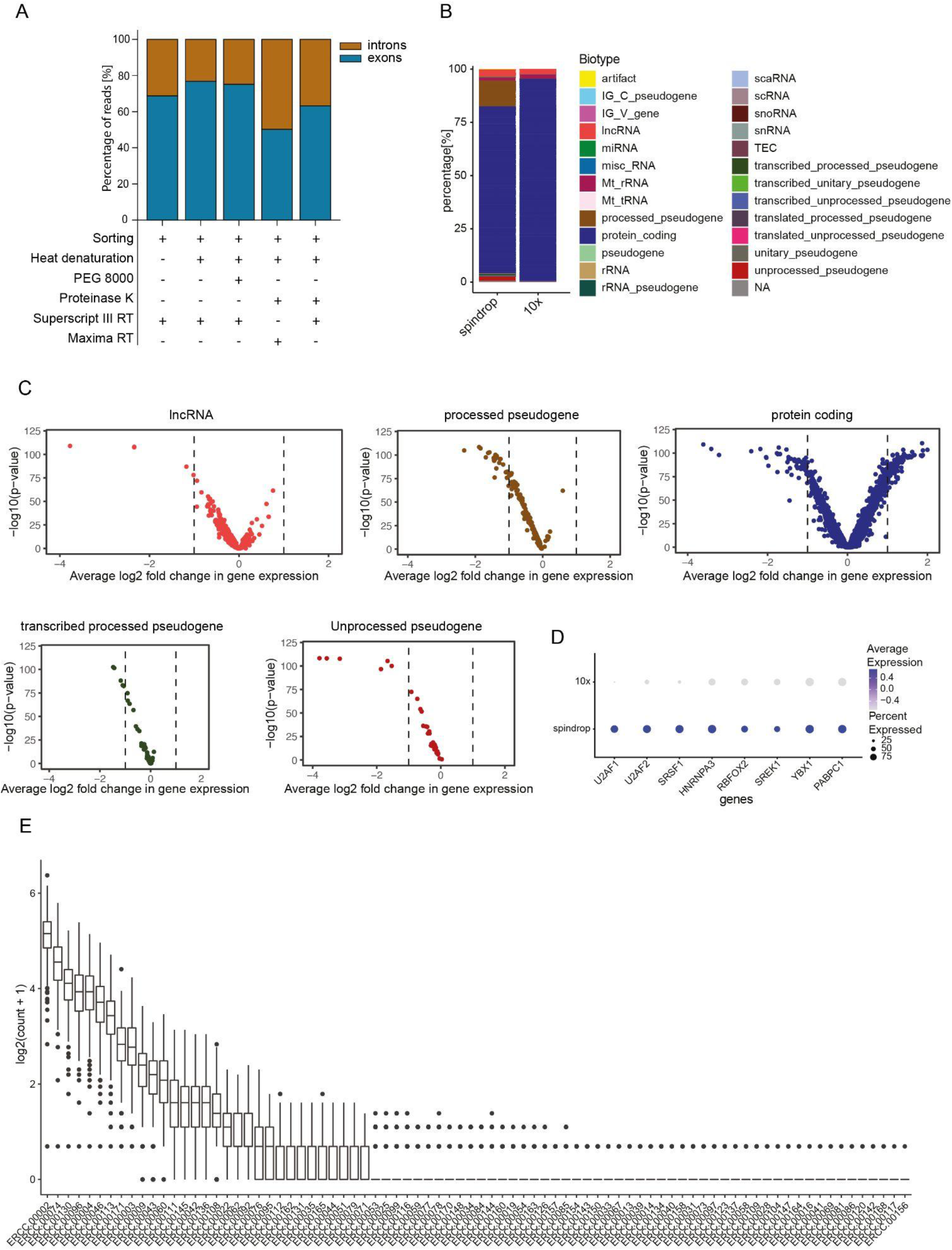
spinDrop biotype repartition and species-mixing assay **A)** Proportional representation of UMI-corrected reads mapping to intronic and exonic regions for different reaction mixtures, showing larger intronic representation with proteinase K and heat denaturation treatment. **B)** RNA biotype proportional representation between 10x and spinDrop HEK293T cells, downsampled to 20,000 reads per cell. **C)** Differential expression analysis using a Wilcoxon rank sum test between HEK293T cells processed using spinDrop (negative log2fold change values) and 10x (positive log2fold change values) for lncRNAs, processed pseudogenes, protein coding genes, transcribed processed pseudogenes and unprocessed pseudogenes. **D)** Differential expression for HEK293T cells between the spinDrop and 10x datasets identifies preferential capture of splicing factors and members of the human spliceosome in the spinDrop dataset, illustrating the potential for spinDrop to uniquely uncover biological processes. **E)** ERCC molecules detected across all sampled droplets from a mouse ES and human HEK293T species-mixing experiment, showcasing the ability of spinDrop to capture spike-ins at a reduced cost through exclusion of empty droplets.

**Extended Data Figure 4.**
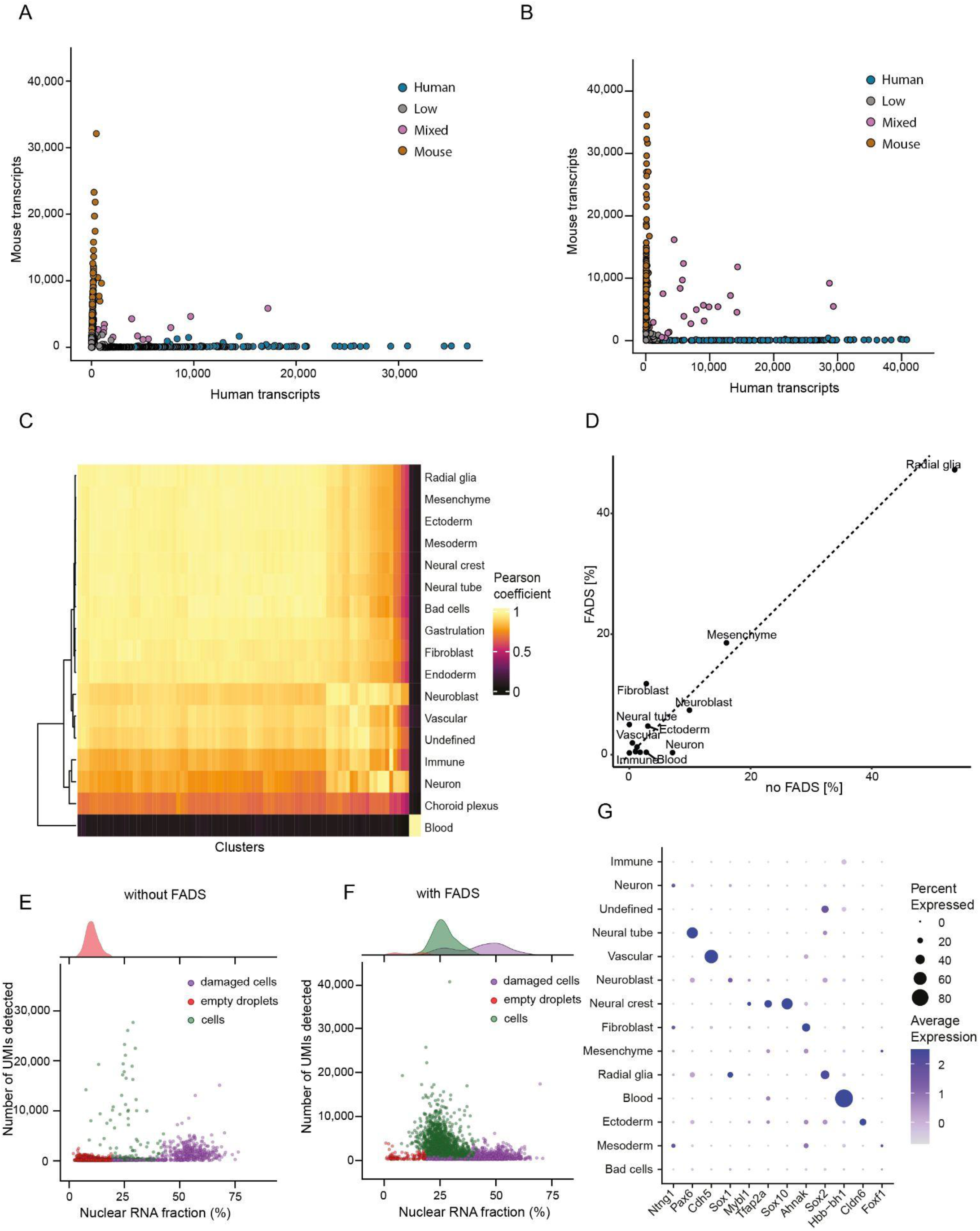
spinDrop enables RNA spike-ins and delivers high-quality data from damaged samples. **A)** Species-mixing Barnyard plot using mES and human HEK293T whole cells, processed with the spinDrop protocol. **B)** Species-mixing Barnyard plot using mES and human HEK293T extracted nuclei, processed with the spinDrop protocol. **C)** Pearson correlation heatmap between the shared embedding clusters and the labelled cell types from La Manno *et al*. using the isolated brain sections of mouse embryos at E10.5. **D)** Proportion of cell types in the spinDrop dataset with and without FADS. **E)** DropletQC quality metrics for the spinDrop embryo dataset without FADS sorting, showing large representation of barcodes for damaged cells and empty droplets. **F)** DropletQC quality metrics for the spinDrop embryo dataset with FADS sorting, showing large representation of barcodes labelled as viable cells. **G)** DotPlot representing core cell type marker expression for different cell types in the spinDrop embryonic dataset.

**Extended Data Figure 5.**
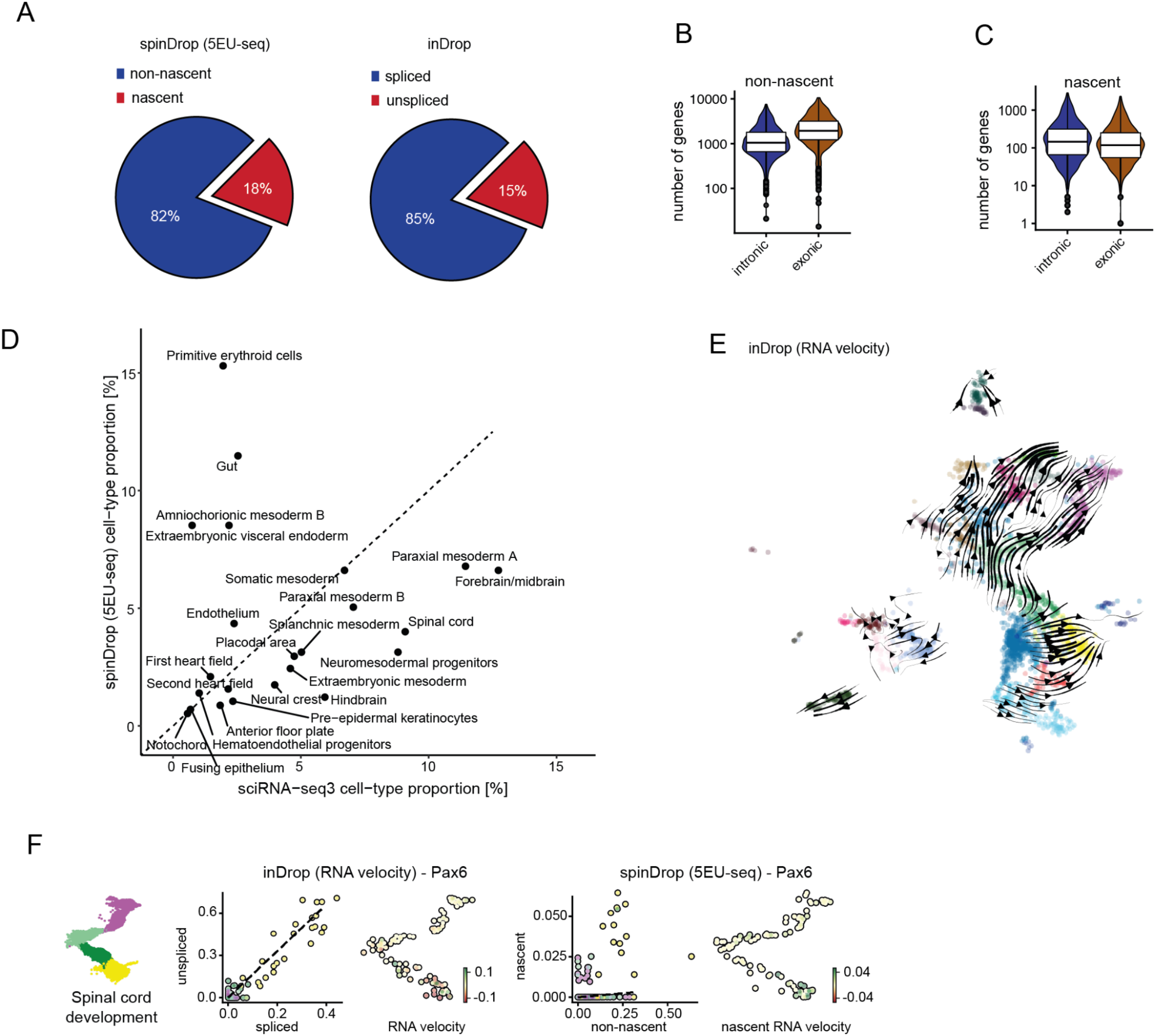
Nascent RNA velocity sequencing using droplet-based 5EU-seq. **A)** Proportion of nascent RNA in the spinDrop 5EU-seq (left) and unspliced read in the E8.5 inDrop mouse organogenesis dataset (right). **B)** Number of genes detected using reads mapping to intronic and exonic regions in the non-nascent RNA 5EU-seq dataset generated using spinDrop. **C)** Number of genes detected using reads mapping to intronic and exonic regions in the nascent RNA 5EU-seq dataset generated using spinDrop. **D)** Proportion of cells for each cell type in the sciRNA-seq3 (x-axis) and 5EU-seq (y-axis) datasets. The 5EU-seq dataset was filtered to barcodes containing reads in both the nascent and non-nascent libraries. **E)**RNA velocity vectors computed using scVelo projected on the inDrop dataset of mouse organogenesis at E8.5. **F)***Pax6* RNA velocity profiles and nascent RNA velocity profiles across the spinal cord developmental trajectory.

## References

1. Svensson, V., Vento-Tormo, R. & Teichmann, S. A. Exponential scaling of single-cell RNA-seq in the past decade. Nat. Protoc. 13, 599–604 (2018).

2. Null, N. et al. The Tabula Sapiens: A multiple-organ, single-cell transcriptomic atlas of humans. Science 376, eabl4896 (2022).

3. Karlsson, M. et al. A single–cell type transcriptomics map of human tissues. Science Advances 7, eabh2169 (2021).

4. Cao, J. et al. A human cell atlas of fetal gene expression. Science 370, (2020).

5. Suo, C. et al. Mapping the developing human immune system across organs. Science 376, eabo0510 (2022).

6. Elmentaite, R. et al. Single-Cell Sequencing of Developing Human Gut Reveals Transcriptional Links to Childhood Crohn’s Disease. Dev. Cell 55, 771–783.e5 (2020).

7. Litviňuková, M. et al. Cells of the adult human heart. Nature 588, 466–472 (2020).

8. Aizarani, N. et al. A human liver cell atlas reveals heterogeneity and epithelial progenitors. Nature 572, 199–204 (2019).

9. Sebé-Pedrós, A. et al. Cnidarian Cell Type Diversity and Regulation Revealed by Whole-Organism Single-Cell RNA-Seq. Cell 173, 1520–1534.e20 (2018).

10. Chari, T. et al. Whole-animal multiplexed single-cell RNA-seq reveals transcriptional shifts across Clytia medusa cell types. Science Advances 7, eabh1683 (2021).

11. Cao, J. et al. The single-cell transcriptional landscape of mammalian organogenesis. Nature 566, 496–502 (2019).

12. Zheng, G. X. Y. et al. Massively parallel digital transcriptional profiling of single cells. Nat. Commun. 8, 1–12 (2017).

13. Tang, F. et al. mRNA-Seq whole-transcriptome analysis of a single cell. Nat. Methods 6, 377–382 (2009).

14. Klein, A. M. et al. Droplet Barcoding for Single-Cell Transcriptomics Applied to Embryonic Stem Cells. Cell 161, 1187–1201 (2015).

15. Macosko, E. Z. et al. Highly Parallel Genome-wide Expression Profiling of Individual Cells Using Nanoliter Droplets. Cell 161, 1202–1214 (2015).

16. Zhang, X. et al. Comparative Analysis of Droplet-Based Ultra-High-Throughput Single-Cell RNA-Seq Systems. Mol. Cell 73, 130–142.e5 (2019).

17. Young, M. D. & Behjati, S. SoupX removes ambient RNA contamination from droplet-based single-cell RNA sequencing data. Gigascience 9, giaa151 (2020).

18. Lun, A. T. L. et al. EmptyDrops: distinguishing cells from empty droplets in droplet-based single-cell RNA sequencing data. Genome Biol. 20, 63 (2019).

19. Muskovic, W. & Powell, J. E. DropletQC: improved identification of empty droplets and damaged cells in single-cell RNA-seq data. Genome Biol. 22, 329 (2021).

20. Staunstrup, N. H. et al. Comparison of electrostatic and mechanical cell sorting with limited starting material. Cytometry A 101, 298–310 (2022).

21. Denisenko, E. et al. Systematic assessment of tissue dissociation and storage biases in single-cell and single-nucleus RNA-seq workflows. Genome Biol. 21, 130 (2020).

22. Hanamsagar, R. et al. An optimized workflow for single-cell transcriptomics and repertoire profiling of purified lymphocytes from clinical samples. Sci. Rep. 10, 1–15 (2020).

23. Yan, F., Zhao, Z. & Simon, L. M. EmptyNN: A neural network based on positive and unlabeled learning to remove cell-free droplets and recover lost cells in scRNA-seq data. Patterns (N Y) 2, 100311 (2021).

24. McGinnis, C. S., Murrow, L. M. & Gartner, Z. J. DoubletFinder: Doublet Detection in Single-Cell RNA Sequencing Data Using Artificial Nearest Neighbors. Cell Syst 8, 329–337.e4 (2019).

25. Wolock, S. L., Lopez, R. & Klein, A. M. Scrublet: Computational Identification of Cell Doublets in Single-Cell Transcriptomic Data. Cell Syst 8, 281–291.e9 (2019).

26. Baret, J.-C. et al. Fluorescence-activated droplet sorting (FADS): efficient microfluidic cell sorting based on enzymatic activity. Lab Chip 9, 1850–1858 (2009).

27. Abate, A. R., Hung, T., Mary, P., Agresti, J. J. & Weitz, D. A. High-throughput injection with microfluidics using picoinjectors. Proc. Natl. Acad. Sci. U. S. A. 107, 19163–19166 (2010).

28. Zilionis, R. et al. Single-cell barcoding and sequencing using droplet microfluidics. Nat. Protoc. 12, 44–73 (2017).

29. Briggs, J. A. et al. The dynamics of gene expression in vertebrate embryogenesis at single-cell resolution. Science 360, (2018).

30. Miles, F. L., Lynch, J. E. & Sikes, R. A. Cell-based assays using calcein acetoxymethyl ester show variation in fluorescence with treatment conditions. J Biol Methods 2, (2015).

31. Sciambi, A. & Abate, A. R. Generating electric fields in PDMS microfluidic devices with salt water electrodes. Lab Chip 14, 2605–2609 (2014).

32. De Rop, F. V. et al. Hydrop enables droplet-based single-cell ATAC-seq and single-cell RNA-seq using dissolvable hydrogel beads. Elife 11, e73971 (2022).

33. Hagemann-Jensen, M. et al. Single-cell RNA counting at allele and isoform resolution using Smart-seq3. Nat. Biotechnol. 38, 708–714 (2020).

34. Bagnoli, J. W. et al. Sensitive and powerful single-cell RNA sequencing using mcSCRB-seq. Nat. Commun. 9, 2937 (2018).

35. Hahaut, V. et al. Fast and highly sensitive full-length single-cell RNA sequencing using FLASH-seq. Nat. Biotechnol. (2022) doi:10.1038/s41587-022-01312-3.

36. Hagemann-Jensen, M., Ziegenhain, C. & Sandberg, R. Scalable single-cell RNA sequencing from full transcripts with Smart-seq3xpress. Nat. Biotechnol. (2022) doi:10.1038/s41587-022-01311-4.

37. Salmen, F. et al. High-throughput total RNA sequencing in single cells using VASA-seq. Nat. Biotechnol. (2022) doi:10.1038/s41587-022-01361-8.

38. La Manno, G. et al. RNA velocity of single cells. Nature 560, 494–498 (2018).

39. Mereu, E. et al. Benchmarking single-cell RNA-sequencing protocols for cell atlas projects. Nat. Biotechnol. 38, 747–755 (2020).

40. Jiang, L. et al. Synthetic spike-in standards for RNA-seq experiments. Genome Res. 21, 1543–1551 (2011).

41. Thomsen, E. R. et al. Fixed single-cell transcriptomic characterization of human radial glial diversity. Nat. Methods 13, 87–93 (2016).

42. O’Flanagan, C. H. et al. Dissociation of solid tumor tissues with cold active protease for single-cell RNA-seq minimizes conserved collagenase-associated stress responses. Genome Biol. 20, 210 (2019).

43. Parekh, S., Ziegenhain, C., Vieth, B., Enard, W. & Hellmann, I. zUMIs - A fast and flexible pipeline to process RNA sequencing data with UMIs. Gigascience 7, giy059 (2018).

44. La Manno, G. et al. Molecular architecture of the developing mouse brain. Nature 596, 92–96 (2021).

45. Stuart, T. et al. Comprehensive Integration of Single-Cell Data. Cell 177, 1888–1902.e21 (2019).

46. Bergsland, M., Werme, M., Malewicz, M., Perlmann, T. & Muhr, J. The establishment of neuronal properties is controlled by Sox4 and Sox11. Genes Dev. 20, 3475–3486 (2006).

47. Knauss, J. L. et al. Long noncoding RNA Sox2ot and transcription factor YY1 co-regulate the differentiation of cortical neural progenitors by repressing Sox2. Cell Death Dis. 9, 799 (2018).

48. Pataskar, A. et al. NeuroD1 reprograms chromatin and transcription factor landscapes to induce the neuronal program. EMBO J. 35, 24–45 (2016).

49. Battich, N. et al. Sequencing metabolically labeled transcripts in single cells reveals mRNA turnover strategies. Science 367, 1151–1156 (2020).

50. Amadei, G. et al. Embryo model completes gastrulation to neurulation and organogenesis. Nature 610, 143–153 (2022).

51. Bergen, V., Lange, M., Peidli, S., Wolf, F. A. & Theis, F. J. Generalizing RNA velocity to transient cell states through dynamical modeling. Nat. Biotechnol. 38, 1408–1414 (2020).

52. Lange, M. et al. CellRank for directed single-cell fate mapping. Nat. Methods 19, 159–170 (2022).

53. Habib, N. et al. Massively parallel single-nucleus RNA-seq with DroNc-seq. Nat. Methods 14, 955–958 (2017).

54. Van Phan, H. et al. High-throughput RNA sequencing of paraformaldehyde-fixed single cells. Nat. Commun. 12, 5636 (2021).

55. Vallejo, A. F. et al. snPATHO-seq: unlocking the FFPE archives for single nucleus RNA profiling. bioRxiv 2022.08.23.505054 (2022) doi:10.1101/2022.08.23.505054.

56. Andrews, T. S., Kiselev, V. Y., McCarthy, D. & Hemberg, M. Tutorial: guidelines for the computational analysis of single-cell RNA sequencing data. Nat. Protoc. 16, 1–9 (2020).

57. Osorio, D. & Cai, J. J. Systematic determination of the mitochondrial proportion in human and mice tissues for single-cell RNA-sequencing data quality control. Bioinformatics 37, 963–967 (2021).

58. Clark, I. C. et al. Targeted Single-Cell RNA and DNA Sequencing With Fluorescence-Activated Droplet Merger. Anal. Chem. 92, 14616–14623 (2020).

59. Zhang, J. Q. et al. Linked optical and gene expression profiling of single cells at high-throughput. Genome Biol. 21, 49 (2020).

60. Bues, J. et al. Deterministic scRNA-seq captures variation in intestinal crypt and organoid composition. Nat. Methods 19, 323–330 (2022).

61. Chen, S., Lake, B. B. & Zhang, K. High-throughput sequencing of the transcriptome and chromatin accessibility in the same cell. Nat. Biotechnol. 37, 1452–1457 (2019).

62. Anagnostidis, V. et al. Deep learning guided image-based droplet sorting for on-demand selection and analysis of single cells and 3D cell cultures. Lab Chip 20, 889–900 (2020).

63. Cole, R. H., Gartner, Z. J. & Abate, A. R. Multicolor Fluorescence Detection for Droplet Microfluidics Using Optical Fibers. J. Vis. Exp. (2016) doi:10.3791/54010.

64. Bedzhov, I. & Zernicka-Goetz, M. Self-organizing properties of mouse pluripotent cells initiate morphogenesis upon implantation. Cell 156, 1032–1044 (2014).

65. Andrews, S. & Others. FastQC: a quality control tool for high throughput sequence data. Preprint at (2010).

66. Galanti, L., Shasha, D. & Gunsalus, K. C. Pheniqs 2.0: accurate, high-performance Bayesian decoding and confidence estimation for combinatorial barcode indexing. BMC Bioinformatics 22, 359 (2021).

67. Dobin, A. et al. STAR: ultrafast universal RNA-seq aligner. Bioinformatics 29, 15–21 (2013).

68. Smedley, D. et al. BioMart – biological queries made easy. BMC Genomics vol. 10 Preprint at https://doi.org/10.1186/1471-2164-10-22 (2009).

